# Epithelial competition determines gene therapy potential to suppress Fanconi Anemia oral cancer risk

**DOI:** 10.1101/2025.02.26.640284

**Authors:** Hunter L. Colegrove, Raymond J. Monnat, Alison F. Feder

**Affiliations:** Department of Genome Sciences, University of Washington, Seattle, WA; Department of Laboratory Medicine and Pathology, University of Washington, Seattle, WA; Department of Bioengineering, University of Washington, Seattle, WA; Herbold Computational Biology Program, Fred Hutch Cancer Center, Seattle, WA

## Abstract

Fanconi Anemia (FA) is a heritable syndrome characterized by DNA damage repair deficits, frequent malformations and a significantly elevated risk of bone marrow failure, leukemia, and mucosal head and neck squamous cell carcinomas (HNSCC). Hematopoietic stem cell gene therapy can prevent marrow failure and lower leukemia risk, but mucosal gene therapy to lower HNSCC risk remains untested. Major knowledge gaps include an incomplete understanding of how rapidly gene-corrected cellular lineages could spread through the oral epithelium, and which delivery parameters are critical for ensuring efficient gene correction. To answer these questions, we extended an agent-based model of the oral epithelium to include the delivery of gene correction *in situ* to FA cells and the competitive dynamics between cellular lineages with and without gene correction. We found that only gene-corrected lineages with substantial proliferative advantages (probability of resisting displacement out of the basal layer ≥ 0. 1) could spread on clinically relevant timelines, and that these lineages were initially at high risk of loss in the generations following correction. Delivering gene correction to many cells minimizes the risk of loss, while delivery to many distinct locations within a tissue maximizes the rate of spread. To determine the impact of mucosal gene therapy in preventing the clonal expansion of pre-cancerous mutations, we compared the expected burden of *TP53* mutations in simulated tissue sections with and without gene correction. We found that when FA cells have elevated genome instability or a *TP53*-dependent proliferative advantage, gene correction can substantially reduce the accumulation of pro-tumorigenic mutations. This model illustrates the power of computational frameworks to identify critical determinants of therapeutic success to enable experimental optimization and support novel and effective gene therapy applications.

**Author summary:** We investigated factors influencing the success of oral mucosal gene therapy for Fanconi Anemia (FA), a genetic syndrome marked by DNA repair defects in conjunction with a heightened risk of cancer. We used a computational model of the oral epithelium to determine how gene therapy corrected cells compete with FA background cells and the best gene delivery approaches to promote effective tissue replacement by gene-corrected cells. We find that gene-corrected cells require strong proliferative advantages to spread effectively, and that initially delivering gene correction to more cells reduces the chance that these cells are stochastically eliminated before they can spread. We also demonstrate that gene correction reduces the accumulation of pro-tumorigenic *TP53* mutations in an FA context, where genomic instability can elevate the mutation rate and FA-specific selective pressures could favor accelerated *TP53* clonal expansion. This research provides a useful framework for guiding mucosal gene therapy experiments and the development of effective oral gene therapy protocols for cancer prevention in FA.

## Introduction

Fanconi Anemia (FA) is an inherited genetic disorder and cancer predisposition syndrome linked to increased risk of bone marrow failure, leukemia, and epithelial cancers. FA arises from biallelic pathogenic mutations in any of 23 *FANC* genes that work in concert as a pathway (the ‘FA pathway’) to detect and coordinate the repair of many types of DNA damage including DNA interstrand crosslinks and double-strand breaks (1–3). This compromised repair capacity is associated with an elevated risk of certain cancers, particularly head and neck squamous cell carcinomas (HNSCC). FA patients experience HNSCC rates several hundred to over a thousand fold greater than the general population (4–8), and a corresponding elevation in mucosal anogenital squamous carcinomas. These cancers often contain pathogenic mutations in *TP53*, a crucial tumor suppressor gene (9–11), and are genomically unstable. Further, they are difficult to treat via standard care chemotherapy and radiation due to the FA-associated constitutional DNA repair defect (8,12–14).

Given these challenges, corrective gene therapy represents a promising strategy for the prevention of FA-associated HNSCC. This strategy has already shown success in FA-associated bone marrow failure, where gene-corrected hematopoietic stem cells (HSCs) can reverse marrow failure and suppress the risk of developing leukemia: HSCs are removed, the specific mutant *FANC* gene is corrected *ex vivo*, and corrected cells are re-implanted into the bone marrow (**Figure 1A**). Recent FA gene therapy clinical studies have found that gene-corrected FA HSCs possess a proliferative advantage over uncorrected FA HSCs (**Figure 1B**, (15–19)). As a result, small numbers of implanted corrected HSCs have been shown to expand over time to reverse FA-associated bone marrow failure.

**Figure 1.**
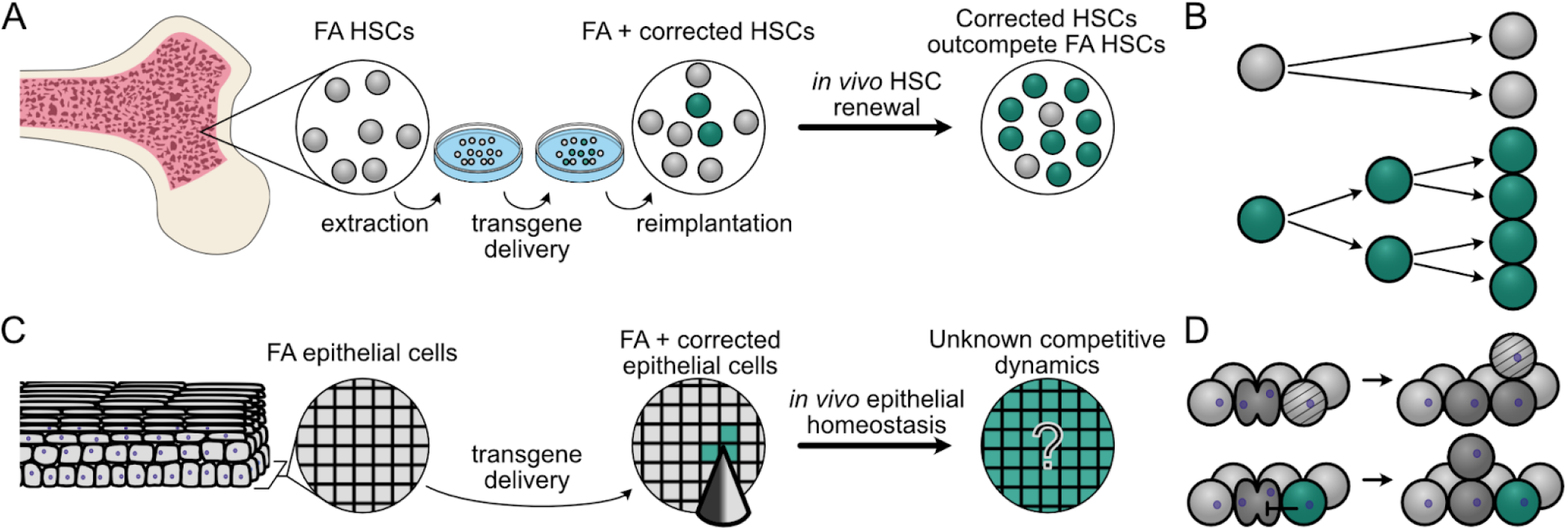
Modeling gene therapy in FA HSCs and oral epithelium. **(A)** Hematopoietic stem cell (HSC) gene therapy in FA. FA HSCs (grey) are harvested from FA patient bone marrow, corrected *ex vivo* (green), then re-implanted to expand *in vivo*. **(B)** Corrected HSCs (green) have a proliferative advantage compared to FA HSCs (grey) and outcompete uncorrected HSCs over time *in vivo*. **(C)** Oral mucosal gene therapy in FA. *In situ* gene delivery (here via microneedles) into oral epithelium to correct the underlying *FANC* gene defect. Over time, corrected cells (green) may or may not clonally expand within FA oral epithelium to replace FA host cells (grey). **(D)** In our model, FA epithelial cells (light grey) are displaced (light grey with bands) by dividing FA cells (dark grey, top), in contrast to gene-corrected epithelial cells (green) that can resist displacement by dividing neighbors (dark grey) to persist and expand in the basal layer proliferative niche (bottom).

The success of HSC gene correction in treating marrow failure suggests that oral gene correction may provide a parallel way to prevent FA-associated HNSCC. However, gene correcting mucosal epithelial cells may prove more challenging than HSCs due to the constraints of tissue architecture. Epithelial tissues are tightly organized, which could limit corrected cell expansion when compared with HSCs in bone marrow (**Figure 1C**). Numerous theoretical and empirical studies have suggested that cells growing under spatial restriction grow more slowly than their unstructured counterparts (20–25). However, cellular lineages can clearly expand within epithelial tissues, e.g., those lineages harboring cancer driver mutations (23,26,27), or spontaneous revertant mutations in the context of heritable skin diseases such as epidermolysis bullosa (28–30). Thus understanding the degree of proliferative advantage required for FA-corrected oral epithelial cells to expand and persist will be essential in devising mucosal gene correction protocols for FA epithelium.

A second major challenge of preventative gene therapy against FA-associated HNSCCs is that oral epithelial cells must be corrected *in situ* rather than removed and reimplanted (**Figure 1C**). This will require identifying efficient mucosal gene correction protocols that favor the expansion and persistence of gene-corrected epithelial cells. One promising new approach is to adapt microneedle arrays originally developed for mucosal vaccination to deliver gene correction reagents (31–36). Microneedle arrays can deliver customizable doses of correction reagents in different spatial configurations, but the optimal array configurations and microneedle properties required to achieve efficient delivery will need to be determined.

Computational modeling provides an efficient framework in which to integrate both different delivery approaches and the effect of epithelial structure on gene correction to optimize therapeutic success.

Agent-based models have a long history of providing valuable insight into cell-cell competition and the emergent spatial dynamics shaping tissue behavior (37–39). We therefore extended an existing agent-based model of squamous epithelium (40) to include gene correction delivered by arrays of diffusible microneedles in order to answer several key questions: 1) what degree of proliferative advantage must corrected cells possess to persist and spread effectively in the oral epithelium; 2) how do different gene correction efficiencies and spatially-structured delivery strategies such as microneedle arrays affect long-term gene correction success; and 3) can mucosal gene correction prevent the clonal spread of pathogenic *TP53* mutations and the progression to oral cancer in individuals with FA? Our results provide a framework for understanding the dynamics of gene therapy in FA oral epithelium, and how best to use mucosal gene correction to reduce cancer risk and improve FA patient health outcomes.

## Results

### *In silico* model of gene correction in the oral epithelia

In order to simulate the impact of gene therapy in epithelial tissues we employed the HomeostaticEpidermis model, a three-dimensional lattice-based hybrid cellular automaton originally developed to model epidermal dynamics over human lifespans (40). In this model, each epithelial cell is an independent agent whose rate of division and death are determined by a diffusible growth factor emanating from the basal layer to simulate a fibroblast source. Dividing cells reside on a basal layer where the resultant progeny are displaced upward, undergo terminal differentiation and are eventually shed at the epithelial surface. This model robustly captures key epithelial features including structure, homeostasis with cell turnover, and realistic clonal expansion of cells that resist death or differentiation. We modified cellular turnover rates to adapt the epidermal model to better recapitulate properties of the oral mucosa (see Discussion and Materials and Methods).

We further adapted the HomeostaticEpidermis model to explore FA oral gene therapy by introducing simulated gene correction to confer a potential proliferative advantage over uncorrected FA cells. In stratified squamous epithelium, clonal expansion may result from a bias in basal layer cell fate decisions that favors the production of progenitor over differentiated cell progeny (41–43). While the mechanisms driving a potential epithelial proliferative advantage in the context of FA gene correction are unknown, evidence from FA HSC correction experiments shows that a selective advantage can be driven by resistance to apoptosis, increased survival in the presence of DNA damaging agents, and longer-term maintenance of self-renewal capacity (1,3,44–47). All of these factors would contribute to longer-term residence of corrected cells in the basal layer, allowing them to retain proliferative capacity. We therefore modeled a proliferative advantage as a persistence coefficient, *pcorr*, which represents the probability that a corrected cell resists displacement out of the basal layer during neighboring cell divisions (**Figure 1D**). This mirrors the approach from Schenck et al. for modeling the proliferative advantage of *NOTCH1* mutations in skin (40), and permitted us to explore the potential effect of gene correction in an FA background, where gene-corrected cells had anywhere from no advantage (or a neutral *p*_*corr*_ = 0) to an extremely strong advantage (*p*_*corr*_ = 1). We first investigated the proliferative advantage necessary to promote reliable expansion of corrected cellular lineages in an FA mucosal background.

### Gene correction requires a strong proliferative advantage to spread over clinically relevant time periods

We hypothesized that gene-corrected cells will require a proliferative advantage compared to uncorrected FA cells to spread through FA oral mucosa. To set baseline expectations for cellular competition without a proliferative advantage, we first considered the case in which a corrected cell has equivalent division behavior to the FA background: that is, when *p*_*corr*_ = 0. We simulated 0.67 mm^2^ (100 × 100 cells) FA tissue sections and delivered gene correction to 10 cells clustered in the tissue center (mirroring single microneedle delivery, **Figure 2A**). Because the median age at which FA patients develop solid tumors is ∼30 years (5,7), we reasoned that corrected cells would need to displace FA cells in the first decade or two after gene correction to be most clinically impactful. We thus followed simulated tissue sections for up to 50 years to determine the fate of corrected cells and their descendants (**Figure 2A**). We considered three fates: all corrected cells could be displaced from the basal layer (i.e., “loss”); corrected cells could overtake 80% or more of the basal layer (i.e., “confluence”); or neither of these outcomes might be reached (i.e., “ongoing” dynamics). Note, our use of the word confluence parallels its standard definition as the percentage of a culture dish covered by a cellular population, as we are interested in the state of gene correction reaching widespread basal layer coverage (a metric for therapeutic success). Under neutral conditions in which corrected cells have no replicative advantage over uncorrected cells (*p*_*corr*_ = 0), nearly all simulations (97%) led to the loss of corrected cells by 50 years (**Figure 2B**) with most corrected cell loss occurring within months of correction (**Figure 2C**). In the remaining 3% of simulations, neutral corrected patches remained, but were of modest size with the largest corrected clone reaching only 0.15 mm^2^. Thus, in the absence of a proliferative advantage, gene correction will not be able to efficiently convert FA epithelium.

**Figure 2.**
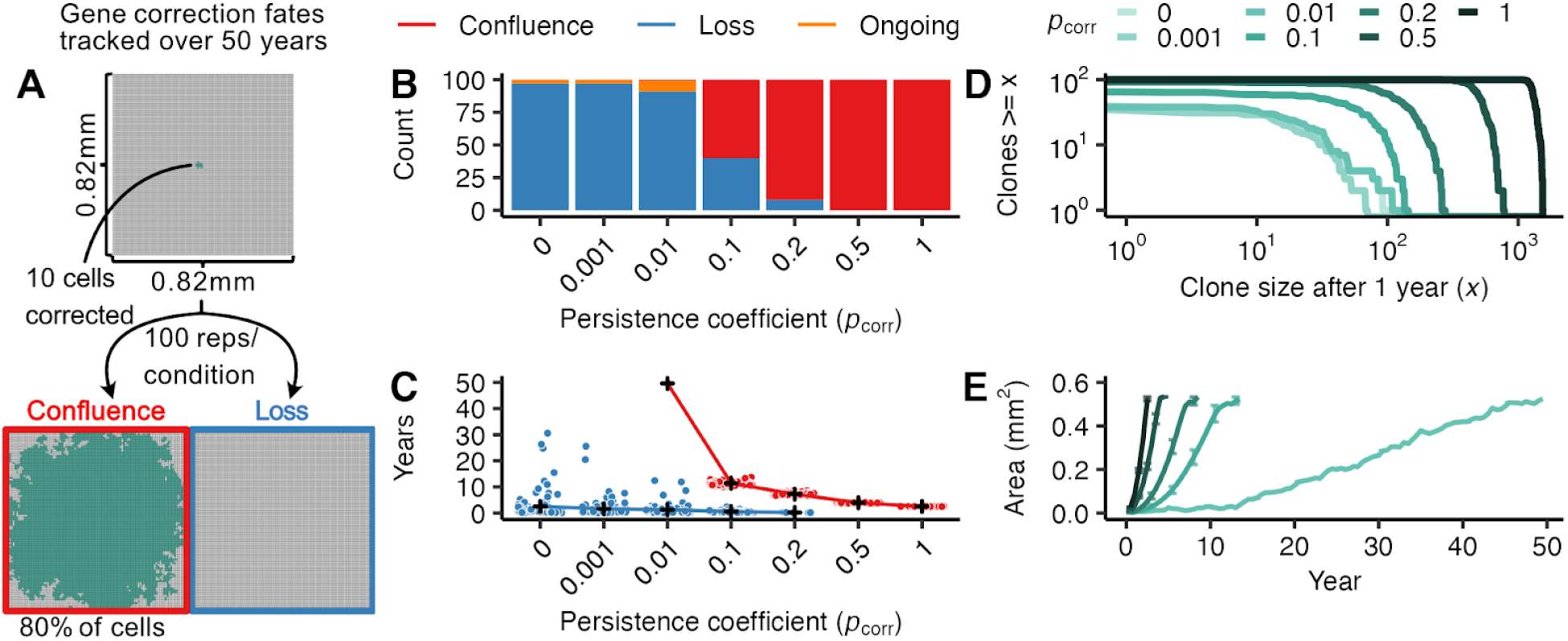
Mucosal gene therapy depends on a strong proliferative advantage of gene-corrected cells. **(A)** 10 gene-corrected cells (green) were tracked over 50 years in a 0.67 mm^2^ FA mucosal tissue section (100 replicate simulations per condition). The gene correction fates were tracked to identify simulations reaching confluence (80% of basal layer cells corrected, red box) or loss (no remaining gene-corrected cells, blue box). **(B)** Number of simulations achieving confluence (red), loss (blue), or ongoing expansion (orange) at 50 years as a function of persistence coefficient (*p* = 0, 0.001, 0.01, 0.2, 0.5, 1). **(C)** Time at which simulations reached confluence or loss as a function of persistence coefficient, with mean times for each *p* _*corr*_ indicated in black (+) and outcome of individual simulations as points. Gene-corrected patches not reaching confluence or loss by 50 years are not plotted. **(D)** Number and size of gene-corrected patches at 1 year as a function of persistence coefficient *p* _*corr*_. **(E)** Average area of confluent gene-corrected patches as a function of time and persistence coefficient *p* _*corr*_. Bars indicate interquartile ranges.

We next investigated how increasing the proliferative advantage might enable more efficient spread of corrected cells through epithelium on clinically-relevant timelines. While we have strong evidence that *FANC* gene reversions possess a proliferative advantage in non-epithelial contexts (e.g., HSCs), we have little direct knowledge about the degree of proliferative advantage conferred by *FANC* gene correction in epithelial tissues. We therefore investigated a wide range of persistence coefficients ranging from unlikely to guaranteed persistence (*p*_*corr*_ = 0. 001, 0. 01, 0. 1, 0. 2, 0. 5, 1) to identify how strong a proliferative advantage must be to ensure durable spread. More biological context for the strength of these persistence coefficients is given in **Supplemental Table 1**. In all experiments, FA cells did not resist displacement (i.e., had a persistence coefficient of 0).

As expected, larger persistence coefficients decreased the probability of corrected cell loss **(Figure 2B)**, with a *p*_*corr*_ ≥ 0.1 required to avoid most loss events. When loss did occur at large persistence coefficients, it occurred early with 88% of losses within the first year after gene correction **(Figure 2C)**. This is analogous to the establishment frequency in population genetics, where once beneficial mutations reach a certain frequency, loss becomes increasingly unlikely. In contrast, gene correction with persistence coefficients closer to neutrality can persist for years or decades before loss though rarely expand to any significant degree. Full clone size distributions one year after correction are shown in **Figure 2D** as a function of *p*_*corr*_.

For gene-corrected cells that escape loss, larger persistence coefficients also increased the speed of corrected cell spread. Corrected cells with smaller persistence coefficients (*p*_*corr*_ < 0. 1) that escaped loss almost never reached confluence in the tissue sections by 50 years. In contrast, gene correction with *p*_*corr*_ ≥ 0. 1 that avoided early loss always reached confluence by 50 years (**Figure 2B**). As the persistence coefficient increased, corrected cell confluence was achieved at progressively earlier times (**Figure 2C**), with *p*_*corr*_ = 1 growing at approximately five times the speed of *p*_*corr*_ = 0. 1 (0.25 ± 0. 027 *SE* mm^2^/year versus 0.048 ± 2. 0 · 10^−3^ *SE* mm^2^/year, **Figure 2E**). These results indicate that effective oral mucosal gene correction must confer a substantial proliferative advantage (*p*_*corr*_ ≥ 0. 1) to ensure persistence and spread on clinically-relevant timescales.

### Optimizing transgene delivery for maximal corrected tissue coverage

Having demonstrated above that individual gene-corrected patches with a strong proliferative advantage could spread locally, we next considered how patch spread was determined by transgene dose and the spatial distribution of corrected cells, where microneedle delivery allows gene dose, spacing, and in-tissue diffusibility to be modified as part of microneedle design and fabrication.

We first investigated how a greater number of corrected cells, equated here to transgene dose, affected the probability of corrected cell loss and time to confluence when delivered in a single compact patch (**Figure 3A)**. As expected, correcting more cells decreased the probability of loss across persistence coefficients (**Figure 3B**), where 30 as opposed to 10 corrected cells decreased the probability of loss for *p*_*corr*_ = 0. 1 from 50% to 10%, but not the time to confluence under favorable conditions (**Figure 3C**). This reflects that the time needed to grow from 3 cells to 30 is short, compared to the longer period needed to replace a majority of cells in a target tissue of size 0.67 mm^2^ (circa 10 years for *p*_*corr*_ = 0. 1). Avoiding early corrected cell loss is the primary advantage conferred by a larger dose at a single injection site.

**Figure 3:**
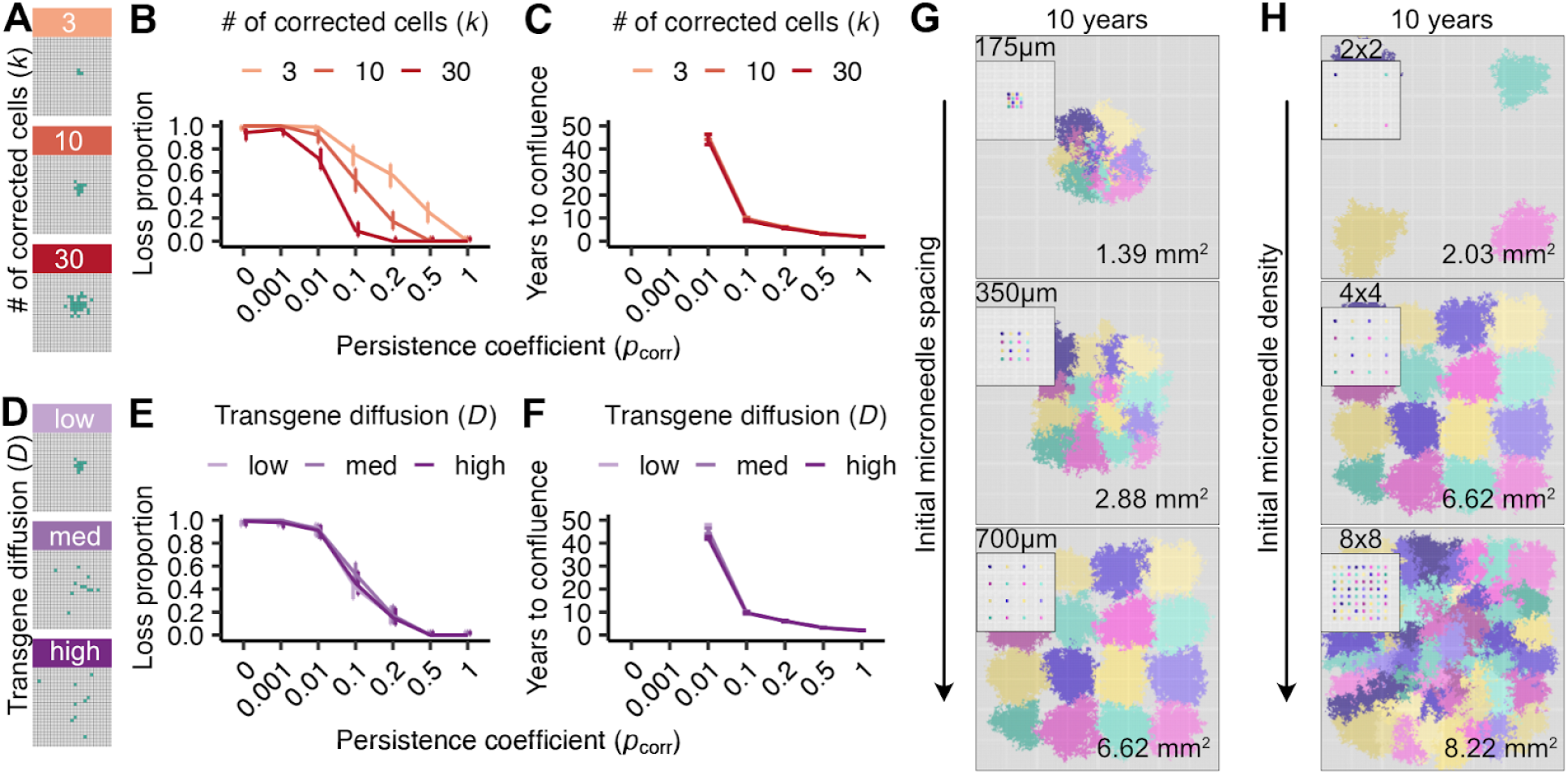
Higher corrected cell numbers and optimal spacing can increase the likelihood of tissue replacement despite clonal interference. Gene-corrected patch loss probability and time to confluence as a function of corrected cell number **(A-C)** and spatial delivery distribution **(D-F)**. 100 simulations/condition were run for each persistence coefficient *p* _*corr*_, with error bars indicating binomial errors **(B,E)** or interquartile ranges **(C, F). (A)** Initial spatial distribution of gene-corrected cells (*k* = 3, 10, 30 cells, in green) with a constant diffusion coefficient (*D* = 2). Probability of gene-corrected patch loss **(B)** and time to confluence **(C)** as a function of persistence coefficient (*p* _*corr*_) and initial corrected cell number (*k*). **(D)** Initial spatial distribution of ten gene-corrected cells (green) as a function of diffusion coefficient (*D* = 2, 10, 20, corresponding to low, medium and high diffusion). Probability of gene-corrected patch loss **(E)** and time to confluence **(F)** as a function of persistence coefficient (*p* _*corr*_) and diffusion coefficient (*D*). Visualizations of gene-corrected cell patches ten years after correction as a function of microneedle spacing **(G)** or density **(H)** on 10.67 mm^2^ tissue sections. Initial arrayed delivery shown in panel insets, and each color represents the descendants of a single microneedle that corrects *k* = 30 cells with *D* = 2. Areas of corrected cell patches at ten years are quantified in the lower right corner of each tissue section.

We next investigated how transgene diffusion from a microneedle injection site might improve corrected cell spread by distributing a fixed number of corrected cells over a larger tissue area. We reasoned that more spatially distributed delivery might allow corrected cells to more effectively compete for space and replace FA cells. We modeled transgene diffusion from a point of delivery with an approximation to Brownian diffusion. The diffusion constant *D* (see Materials and Methods) was examined for *D* ∈ (2, 10, 20) to capture low, intermediate and high degrees of spatial diffusion from 10 initial corrected cells (**Figure 3D**). Surprisingly, increased diffusibility did not affect the probability of corrected patch loss or its rate of spread (**Figure 3EF**). One explanation is that early stochastic loss of individual corrected cells is more likely driven by being displaced by neighboring cells (regardless of correction status), rather than a failure to displace corrected neighboring cells when dividing. These results indicate that efforts to enable wider transgene diffusion from single microneedles is unlikely to improve the spread of corrected cell patches.

A key advantage of microneedles is their ability to be arrayed in different configurations to optimize transgene delivery. In light of finding that transgene diffusibility minimally affected the rate of spread from single needles, we investigated how microneedle arrays could be used to optimize transgene delivery and tissue correction over a 10 year period.

We first asked how needle spacing might be configured to avoid competition and promote spread within a tissue section. We reasoned that individual microneedles could be designed to minimize loss, with interference between microneedle sites a potentially more important determinant of the spread of corrected cells as illustrated in **Figures 3D-F**. We investigated the impact of needle spacing on transgene delivery, corrected cell persistence and rate of spread using a 4×4 array of 16 low diffusibility (*D* = 2) microneedles in which each needle corrected 30 cells to confer a strong proliferative advantage (*p*_*corr*_ = 0. 1). The needle spacings we explored, of 175 µm, 350 µm, and 700 µm, reflect commonly engineered interneedle distances, where our target for gene correction and spread was a 10.67 mm^2^ tissue section (**Figure 3G**). We found that the most widely spaced microneedle arrays corrected more than five times as much tissue area in a 10 year period than the most tightly spaced arrays for a given number of microneedles. In tightly spaced arrays, corrected cells at the perimeter expanded more than those in the interior; this apparent competition among corrected cells was not observed in the most widely spaced arrays. Thus appropriate transgene delivery spacing has the potential to minimize interference or competition of corrected cells and maximize the area of corrected tissue.

Microneedle arrays can be engineered with substantially higher densities than the ones we examined above. Although tightly spaced arrays led to a slower rate of confluence, they achieved greater localized tissue coverage than did widely spaced arrays **(Figure 3G)**. We found that further increasing needle densities led to more complete conversion of a given tissue area, but did not limit the approach to confluence **(Figure 3H)**. For example, increasing microneedle density to 8×8 on a fixed backing size of 4.43 mm^2^ increased tissue correction by ∼25%, whereas decreasing microneedle density to 2×2 on the same backing patch size decreased the tissue correction by ∼70%. These results indicate that dense microneedle array transgene delivery in conjunction with a strong proliferative advantage can provide widespread tissue level correction even when promoting potential spatial competition among corrected cells and clones. Thus our model should be useful for testing a broad array of potential gene correction arrays to optimize gene delivery under different clinical constraints (see Discussion).

### Gene correction reduces *TP53*^*–*^ clonal expansion in genomically unstable FA epithelium

If FA gene-corrected cells can spread through an FA oral epithelium, how might this counter HNSCC risk or progression? An early stage in the development of FA-associated and sporadic HNSCCs is the clonal expansion of pathogenic mucosal *TP53* mutations (9–11). We thus asked if FA gene correction could counter the spread of *TP53* mutations in oral epithelium. In our model, mutations in a single copy of *TP53* can occur in either corrected (here denoted *FANC*^*+*^) or FA (here denoted *FANC*^*–*^) cells according to mutation rates 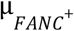 and 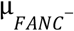 (**Figure 4A**), with *TP53* mutations conferring persistence coefficient advantages of 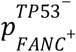 and 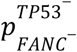, respectively. As before, corrected *FANC*^*+*^ cells without *TP53* mutations have a persistence coefficient of *p* _*corr*_, so corrected *FANC*^*+*^ cells with a *TP53* mutation have a total persistence coefficient of 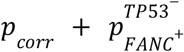 (**Figure 4B**). We estimated a mutation rate 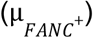 and persistence coefficient 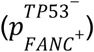 of *TP53* mutations in a *FANC*^*+*^ background based on the rate of occurrence and size of *TP53* clones in healthy human esophageal tissue (see Materials and Methods). In the absence of direct measures, we initially assumed that the rates of occurrence and clonal expansion of *TP53* mutations were similar in *FANC*^*+*^ and *FANC*^−^ tissues (i.e., 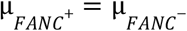 and 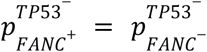). We reasoned that even if FA gene-corrected and uncorrected cells produced the same number of *TP53* mutations with comparable proliferative advantages, gene correction might make *TP53* mutant cells less able to displace *FANC*^*+*^ as opposed to *FANC*^−^ cells, thus restricting their expansion potential.

**Figure 4.**
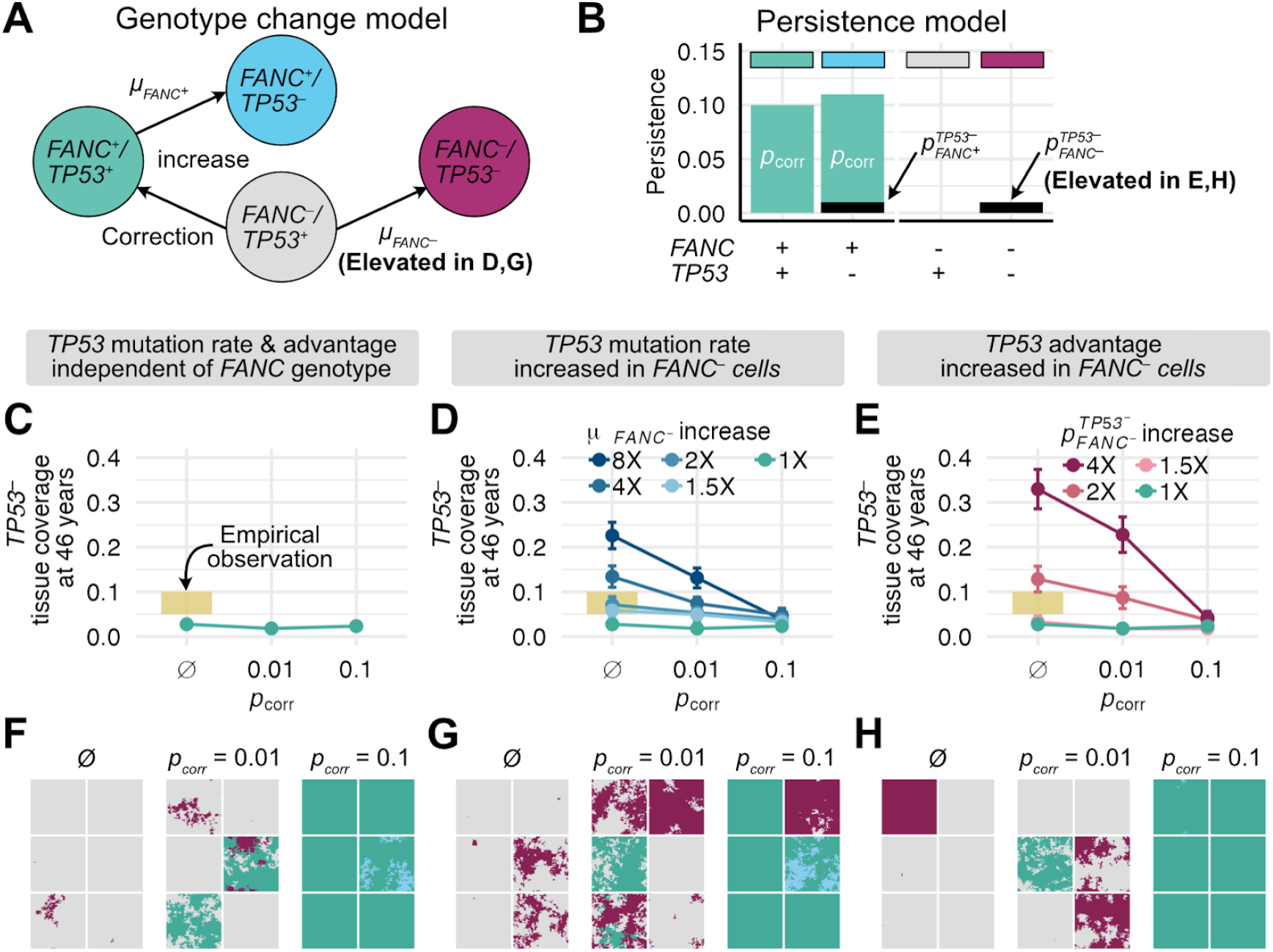
Gene correction reduces spread of *TP53* mutations through FA tissue. **(A)** Model of four cellular genotypes at two loci: FA cells (*FANC*^−^/*TP53*^+^) in grey; FA cells with *TP53* mutations (*FANC*^−^/*TP53*^−^) in magenta; gene-corrected cells (*FANC*^+^/*TP53*^+^) in green; gene-corrected cells with *TP53* mutations (*FANC*^+^/*TP53*^−^) in cyan. Arrows indicate transitions that can occur via mutation or gene correction. **(B)** Genotype-specific persistence coefficients for **(A). (C-H)** *TP53*^−^ tissue coverage as a function of gene correction at 46 years in 0.33 mm^2^ simulated tissue sections, comparing no correction (ø) to correction with *p* _*corr*_ = 0. 01 or *p* _*corr*_ = 0. 1. Panels used 100 simulations at a given persistence coefficient except in panels **C,F** where 300 simulations were performed. **C-E** show the proportion of tissue coverage at 46 years with **(C)** comparable *TP53* mutation rates and persistence coefficients regardless of FA genotype, **(D)** an elevated *TP53* mutation rate in *FANC*^−^ cells 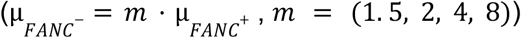 and equivalent *TP53* persistence coefficients or **(E)** equal *TP53* mutation rates in *FANC*^−^ and *FANC*^+^ cells with elevated *TP53* persistence coefficients in *FANC*^−^ cells 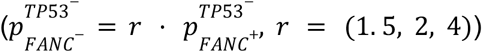. Error bars represent standard errors. Yellow bars represent *TP53* tissue coverage range from Martincorena et al. (48) for normal esophagus. **(F-H):** six representative tissue sections at 46 years from **C-E**, with colors corresponding to genotypes shown in **(A). (G)** consequence of of an 8-fold increase in 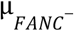, and **(H)** a 4-fold increase in 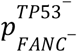.

To test this idea, we asked if gene correction could limit *TP53* mutational prevalence in FA tissue when *TP53* mutation rates and proliferative advantages were comparable for *FANC*^−^ and *FANC*^*+*^ cells (**Figure 4CF**).

We simulated 0.33 mm^2^ (70×70 cells) tissue sections in which a single microneedle gene corrected 30 cells (*D* = 2) with persistence coefficients of *p* _*corr*_ = 0. 01 or *p* _*corr*_ = 0. 1 at the tissue center, then followed these sections up to 50 years. We compared these corrected tissue sections to uncorrected tissue section controls, and also plotted *TP53*^−^ expansion in healthy esophageal tissue as a reference (48). We found that *TP53* mutations reached similar tissue frequencies over 46 years regardless of gene correction status (**Figure 4CF**), as might be expected under an additive fitness model. Gene correction restricted *TP53*^−^ spread in *FANC*^−^ cells, while mutations still arose and expanded within corrected *FANC*^*+*^ cells leading to similar overall tissue frequencies (compare cyan *FANC*^+^/*TP53*^−^ cells with *p* _*corr*_ = 0. 1 in **Figure 4F** tissue sections to uncorrected magenta *FANC*^−^/*TP53*^−^ cells). Gene correction might be beneficial if *FANC*^−^/*TP53*^−^ double mutants are more likely to progress to cancer than *FANC*^*+*^/*TP53*^−^ mutants. However, this is unlikely to lower the overall *TP53*^−^ burden if *TP53* mutation rates and proliferative advantages are comparable between corrected and uncorrected tissue. Full 46-year tissue coverage trajectories from these simulations are shown in **Supplemental Figure 1A-C**, and demonstrate that *TP53*^−^ tissue coverage over time is similar for each value of *p* _*corr*_.

The above results assumed that the *TP53* mutation rate and mutant proliferative advantage are identical regardless of *FANC* genotype. However, the dramatically increased risk of HNSCC in FA individuals likely reflects genomic instability in FA that could generate and promote the spread of pathogenic mutations through FA epithelia. We therefore investigated how FA genotype affects the rate of *TP53* mutation generation and spread, and thus could determine the outcome of FA gene correction.

We first considered that *FANC*^−^ cells have a higher mutation rate than *FANC*^*+*^ cells (49–51). As there have been a range of values reported for this increase (up to 8-30x), we reran the experiments above to determine outcomes with *TP53* mutation rates in uncorrected *FANC*^−^ cells 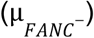 of 1.5-, 2-, 4-, or 8-fold higher than in corrected *FANC*^*+*^ cells (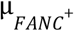, **Figure 4DG**). In the absence of FA gene correction, 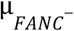 increased by a factor of eight led to a corresponding increase in the tissue burden of *TP53*^−^ cells to cover 23% of tissue sections at 46 years, or nearly 10-fold higher than the coverage estimate of 2.7% under the background mutation rate (**Figure 4D**). While elevated mutation rates produced more *TP53*^−^ clones, individual clone sizes remained similar to those under baseline mutation rates (**Supplemental Figure 2AB**). We found that by introducing gene correction with a small persistence coefficient (*p*_*corr*_ = 0. 01, **Figure 4D**), *TP53*^−^ tissue coverage was modestly reduced compared to uncorrected tissue (a reduction of 42% in simulations where *FANC*^−^ cells had a 8x elevated mutation rate). In contrast, gene correction with a larger persistence coefficient (*p*_*corr*_ = 0. 1, **Figure 4D**) resulted in tissue sections with almost no additional *TP53* mutations above baseline simulations that used background mutation rates. This effect depended on gene-corrected *FANC*^*+*^ cells replacing more mutable *FANC*^−^ cells: in the 2% of *p*_*corr*_ = 0. 1 simulations in which gene corrected cells were stochastically lost, *FANC*^−^*/TP53*^−^ cells expanded as if they were in uncorrected tissue (see panel inset in **Figure 4G**). Full 46-year tissue coverage trajectories from these simulations are shown in **Supplemental Figure 1D-F**.

We next considered that *TP53* mutations might confer a stronger proliferative advantage in FA cells than in gene-corrected cells (i.e., 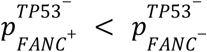). The basis of this assumption is observed non-additive effects on the rate of cell turnover and tumor formation in cells harboring both *TP53* mutations and FA mutations (see Discussion, (52–55)). We parameterized possible fitness advantages using human lymphoblastoid and mouse HSC experimental data (see Materials and Methods). To test the impact of this effect, we reran the experiments above with identical *FANC*^−^ and *FANC*^*+*^ mutation rates, but with 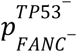 increased by a factor of 1.5-, 2-, or 4-fold compared to 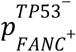 (**Figure 4EH**). In the absence of gene correction, when 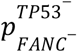 was increased by a factor of 4, there was a corresponding increase in the tissue burden of *TP53* ^*-*^ cells to cover 33% of tissue sections at 46 years, compared with 2.7% under the background proliferative advantage. In contrast to the mutation rate experiments, increased *TP53*^−^ coverage was driven primarily by larger clones in conjunction with a more modest increase in clone number (**Supplemental Figure 2CD**). As was the case for gene correction affecting mutation rates, gene correction with a small persistence coefficient (*p*_*corr*_ = 0. 01, **Figure 4E**) modestly reduced the *TP53*^−^ tissue burden by ∼31% among cells with a 4x elevated proliferative advantage compared to uncorrected tissue. Gene correction with a large persistence coefficient (*p*_*corr*_ = 0. 1, **Figure 4E**) resulted in tissue sections with almost no excess *TP53*^−^ mutational coverage beyond baseline simulations in which *FANC*^−^ and *FANC*^*+*^ cells have identical persistence coefficients associated with *TP53* loss. Again, the rapid spread of gene-corrected cells largely abrogated the proliferative advantage of *TP53* mutations in a *FANC*^−^ background. Full 46-year tissue coverage trajectories are shown in **Supplemental Figure 1G-I**. These results collectively indicate that FA-directed mucosal gene therapy has the potential to reduce *TP53*-associated HNSCCs in FA, and that likely mechanisms include a suppression of FA-associated genomic instability and the strong selective pressure for loss of *TP53* function.

## Discussion

We extended an agent-based model of squamous epithelial clonal competitive dynamics (40) to study epithelial gene correction in oral mucosa. We chose this approach over the more analytically-tractable branching-process models widely used to study tumorigenesis (56,57) because it permitted us to more faithfully capture biologically important spatial constraints on the spread of mutations and gene correction in an epithelial context. These constraints are critical to understand in order to develop highly effective gene correction protocols for FA or other epithelial diseases, as spatial and non-spatial growth patterns have well-characterized diverging dynamics (20–25). This approach recapitulated certain results well-appreciated from branching process models (i.e., correcting more cells limits stochastic loss, cells with small fitness advantages can persist at low frequencies at length), but it also provided practical observations into potential spread rates of corrected patches, optimal strategies for designing correction arrays (see more below), and the role of interference both among patches and with pathogenic clones, all of which rely on incorporating spatial competition explicitly.

A compelling biological rationale for extending the HomeostaticEpidermis (HE) model is the many shared features of oral mucosa and epidermis. These tissues are contiguous at the proximal end of the GI tract, and share common embryological origins, many biological features and pathogenic mechanisms. Alterations in progenitor cell dynamics in these epithelia can promote both pre-neoplastic and neoplastic disease by facilitating the expansion of mutant clones (23,26,27,58–60). We modified the HE model to incorporate oral mucosa-specific data where they exist (e.g., on oral mucosal turnover, cellular density) and further note that many epidermis-specific features (e.g., keratinization, presence of hair follicles, certain wound healing properties) are not represented in the HE model and thus do not impact our findings.

Implementation of our adapted HE model provided several important insights into oral mucosal dynamics, and the potential to counter cancer risk in individuals with FA: 1) gene correction needs to confer a strong proliferative advantage to escape stochastic corrected cell loss, and to promote tissue replacement on clinically-relevant decadal timescales; 2) tissue replacement success can be maximized by increasing delivery dose and by distributing tissue delivery spacing to speed tissue replacement while reducing competitive interference among corrected lineages; and 3) gene correction has the substantial potential to reduce the high risk of HNSCC in FA if it lowers the mutation rate and/or proliferative advantage of oncogenic mutations. Each of these results and their translational importance are discussed in greater detail below.

We found that gene corrected cells will require a strong proliferative advantage (i.e., *p*_*corr*_ ≥ 0. 1) and rapid spread within the oral epithelium for effective FA oral gene correction. Whether FA gene correction can confer a strong proliferative advantage has been most convincingly demonstrated in the hematopoietic system (15–19), and is plausible though less well-documented in oral mucosa. The potential for persistence and rapid spread in related squamous epithelia is exemplified by heritable skin diseases such as epidermolysis bullosa, where patients may display mutationally revertant skin patches 2 cm^2^ or larger, even at ages as young as 8, and with geometries inconsistent with emergence during development (28–30). Similar data exist from individuals with ‘ichthyosis-with-confetti’, who have large revertant skin patches caused by mitotic recombination that can reach up to 4 cm^2^ in size and have been documented in individuals as young as 18 (61). Comparably rapid tissue replacement may occur in the oral mucosa, which has faster measured cell turnover than does epidermis (41,62–64). If FA gene correction confers only a weak intrinsic mucosal fitness advantage, this might be augmented by engineering additional genetic alterations. For example, recent studies have found that *NOTCH1* mutations can confer a fitness advantage that promotes clonal expansion while minimally affecting cancer risk (26).

Our model predicts that gene correction will be most beneficial at suppressing cancer risk in FA if it reduces genomic instability or counters the spread of pathogenic epithelial mutations. There is a strong biological rationale for these suppositions. FA cells accumulate chromosomal aberrations due to a DNA damage sensing and repair deficit (9,65–67), and tumors that arise on this background are further mutated and chromosomally rearranged (9). Complementing the FA defect suppresses these phenotypes, whether by gene correction (15–19) or spontaneous genetic reversion (68–74), in FA patient HSCs. The restoration of FA function reduces DNA damage markers, restores chemoresistance and suppresses genomic instability. Moreover, FA gene correction also has the potential to suppress the spread of pathogenic mutations and tumor progression. The plausibility of these additional effects has been observed in *Fancd2*/*Tp53, Fancc/Tp53* and *Fanca*/*Tp53* double knock-out mice: all have significantly lower rates of cancer-free survival than either single mutant or control mice for epithelial as well as non-epithelial tumors (54,55,75). The importance of the FA-*TP53* interaction is illustrated in mouse HSCs, where *Tp53* elimination on a *Fanca* mutant background increases cellular turnover rates to a greater extent than on a *Fanca*-competent background (53). A potential mechanism for this increased fitness advantage is that persistent DNA damage in FA activates *TP53* to arrest cell cycling for DNA damage repair (52,76). *TP53* loss, in contrast, confers a growth advantage by suppressing proliferation arrest to allow continued cell division in genomically unstable or mutant cells (52,53,77). Gene therapy to restore FA pathway function could thus suppress multiple FA-dependent mechanisms that promote genomic instability, and that provide strong selective pressure for the emergence of *TP53* mutations.

Our oral mucosal epithelial agent-based model could be extended or improved in several ways. First, we could more explicitly model the basal layer stem cell hierarchy. This would likely improve the biological accuracy of the model without substantially altering global tissue dynamics (40). Second, our use of a persistence coefficient could be improved using experimental data that quantify the survival benefits of gene correction in an FA background, when such data become available in an epithelial context. In FA-corrected HSCs, proliferative advantage appears to reflect a combination of reduced apoptosis, increased survival after DNA damage, and/or improved self-renewal capabilities (44–46). Better characterization of these potential mechanisms in FA epithelia should improve modeling insight and predictive outcomes.

More sophisticated genetic models of FA or other diseases could also be developed to provide additional mechanistic insight while capturing between-individual differences. For example, we explored only the effects of single deleterious *TP53* mutations as some *TP53* mutations exhibit dominant-negative effects with heterozygous clonal expansion in mice (27,78,79). A second way to more accurately model FA would be to incorporate loss of function of one of the 23 FA genes with loss of function variants in *ADH5 and ALDH2*. These two genes are key members of aldehyde detoxification pathways that limit the toxicity of formaldehyde and acetaldehyde, both common and unavoidable sources of endogenous and exogenous DNA damage that activates *TP53*-dependent DNA damage signaling and FA pathway activity (80). Our modeling approach can be extended to predict a range of fitness effects of these inputs in both FA and FA gene-corrected cells. A final note is that our modeling framework can accommodate a wide range of new data from many gene delivery platforms in addition to microneedle-mediated gene delivery (31–36).

Gene therapy represents a promising approach to treat or prevent a growing number of disease states and predispositions. Computational modeling, as illustrated here, can provide a useful framework that identifies important determinants of therapeutic success across many diseases and their target cells, tissues or organs. A critical next step in determining the potential of FA oral epithelial gene therapy is to quantify the proliferative advantage of gene-corrected cells in FA epithelia. It may be possible to estimate proliferative advantages now for a subset of FA patients using deep sequencing of oral mucosal biopsies, and from these data estimate reversion rates and the likely revertant proliferative advantage for FA-revertant cells. Several new mouse models of FA (see, e.g., (75)) should soon allow modeling predictions to be tested directly by lineage tracking to quantify the fitness effect of FA gene correction. These mouse models have the advantage of adding spatial distribution data on the spread of FA-corrected cells. All of these approaches should provide new insight into the competitive dynamics of FA gene-corrected cells.

A final important point is that many gene therapy or editing protocols are now focused on *in vivo* gene delivery, where success will depend on integrating target cell and tissue type-specific features with delivery method-specific details (81,82). Our approach allows many of these protocol-, patient- and disease-specific details to be captured and explored *in silico*, and to aid the design of clinical protocols with the highest likelihood of success (83). Modeling also has the potential to minimize excessive animal testing and patient sampling. Key protocol details (delivery vehicle, delivery route, dose and timeline) can be identified and then tracked to measure protocol effectiveness. Clinical data from small *n* trials - or even single patients - can then be used to improve protocol design, implementation and tracking of patient-specific outcomes (see, e.g., the recent *n* = 1 *in vivo* editing of CPS1 deficiency (84)). All of these efforts should substantially improve the design, implementation and eventual success of gene therapy protocols to treat or prevent a wide range of human diseases.

## Acknowledgements

We acknowledge useful discussion with Kim Woodrow, Craig Dorrell, Markus Grompe, members of the Feder lab at UW and helpful comments from three anonymous reviewers. This work was supported by a Research Award from the Fanconi Cancer Foundation to RJM and by 1DP2CA280623 to AFF.

## Materials and methods

### Overview of the model

#### Model Framework

We extended a cellular automaton model of epithelial cell division and homeostasis in the Hybrid Automata Library (85), HomeostaticEpidermis (HE) (40), to simulate the dynamics of oral epithelial cells and corrective gene therapy. Briefly, HE simulates cells as independent agents on a three dimensional lattice that divide and die according to the available concentration of a growth factor diffusing from the basal layer (see for full parameterization of this growth factor gradient). During a cellular division event, one daughter cell occupies the location of the parent cell, while the other is positioned either vertically above or in one of the four orthogonal neighboring positions within the basal layer Von Neumann neighborhood. The placement of this second daughter cell is governed by the division location probability *ω*, where ω = 0. 2518617 represents the probability of vertical placement, and each orthogonal direction has a probability of (1 − ω)/4. If a daughter cell occupies a neighboring position within the basal layer, the displaced cell stratifies out of the basal layer and all cells above it are displaced upwards. The epithelial dynamics of this study focus only on the basal layer, as once a cell stratifies to the suprabasal layers, cells do not de-differentiate and re-enter the basal layer. Cells can mutate during division, which we examine in the latter half of our study. This model reproduces key behaviors of epithelial tissues, including the distribution of mutated clone sizes over time. As such, most existing model parameters are kept at their default values to preserve homeostatic tissue function. New functions and parameters specific to our model are described below.

In our model, all cells begin with mutant *FANC* gene function (i.e., an FA phenotype), as expected in an inherited disorder. Throughout most of the analyses, this FA phenotype does not affect the normal maintenance of homeostatic equilibrium or cellular behavior, although in section *Lineage competition between gene-corrected cells and TP53 mutants*, we describe how uncorrected FA cells may affect genome stability and control of *TP53*^−^ mediated clonal expansions.

#### Adaptation of epidermal model for the oral epithelium

We sought to preserve homeostatic behavior of the HE model while accounting for the increased turnover rate of oral epithelial cells compared to epidermal cells. Under the default parameters in the HE model, basal cells divide on average 0.4 times per week (**Supplemental Figure 3**). However, basal cells in the oral epithelium and esophagus divide approximately twice per week (27,41,62). We therefore rescaled time in our model such that each reported year of simulation time in the oral epithelium corresponds to 4.5 years of simulation time in the HE model.

To reflect the higher density of cells in the oral epithelium versus epidermal tissue, tissue section sizes are determined using a density of 15,000 basal layer cells/mm^2^ (23,41), instead of 10,000 basal layer cells/mm^2^ in the HE model (40,86).

#### Gene Correction

To mimic *in vivo* microneedle gene delivery that corrects FA oral epithelial cells, we allowed FA cells to be transformed into corrected cells by exposure to a simulated transgene at a pre-specified treatment time. We apply the *in silico* transgene delivery to the center of simulated tissue sections except when otherwise noted at the start of the simulation. Transgene spread through the tissue is governed by an approximation to Brownian diffusion away from the injection site. Specifically, for each cellular position on the tissue basal layer, we determine the relative probability of correction by integrating the PDF of a bivariate Gaussian distribution over the cell’s x and y coordinate boundaries on the lattice. We then sample *k* cell coordinates for correction from this probability distribution without replacement. To simulate a transgene injection with greater or lesser diffusion, we examine Gaussian distributions with different variances (*D*) centered on the injection site coordinates (*x* _*inj*_, *y* _*inj*_) as follows:

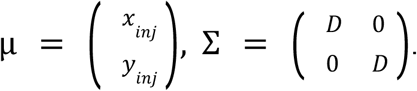

Corrected cells can have altered dynamics in the basal layer that may allow them to spread, in analog to the ability of gene-corrected cells to spread in the hematopoietic compartment. In stratified squamous epithelium, clonal expansion occurs due to an imbalance in the fate of progenitor cells in the basal layer

(23,41,43). We model this mechanism via a persistence coefficient (*p* _*corr*_), which prevents a cell from being displaced by its neighbor cell’s division. Specifically, when a cell divides and randomly chooses to displace a neighboring cell possessing a persistence coefficient *p* _*corr*_, the neighboring cell will not be displaced vertically with probability *p* _*corr*_, and will be displaced vertically with probability *1 - p* _*corr*_. If the neighboring cell is not displaced, the dividing cell instead places its daughter cell vertically and does not displace any neighboring cells. This mirrors the implementation of *NOTCH1* mutations in Schenck et al. (40). FA cells are assumed to have *p* _*corr*_ = 0 throughout.

### Interrogating clonal expansion of corrected cells in an FA background

To investigate the spread of gene correction through the oral epithelium, we simulated FA tissue sections of size 0.67 mm^2^ (100 cells x 100 cells). We introduced a single corrected patch of *k* cells with persistence coefficient *p*_*corr*_ in the center of the basal layer of the tissue as described above and recorded tissue states longitudinally for up to 50 years at six month intervals. Unless otherwise noted, *k = 10* and *D* = 2, reflecting a tightly clustered patch of few corrected cells. Simulations were stopped if no corrected cells remained in the basal layer (‘loss’) or if the corrected patch of cells occupied 80% of the basal layer (‘confluence’). Simulations reaching neither endpoint in 50 years were designated as ‘ongoing’. We ran 100 correction replicates for *pcorr* = (0, 0. 001, 0. 01, 0. 1, 0. 2, 0. 5, 1. 0) and recorded the proportion of simulations in which correction was lost or reached confluence and the time at which that event occurred. We repeated these analyses examining different numbers of corrected cells (*k* = 3, 10, 30) and different degrees of transgene diffusion (*D* = 2, 10, 20). Clone size distributions were determined by computing the number of cells associated with each of the corrected patches for each persistence coefficient after one year. To determine the expansion rates of corrected patches, we examined only simulation replicates reaching confluence and converted the number of corrected cells at each timepoint (six-month intervals) into area assuming each cell was 66.7 µm^2^ (23,41).

### Lineage competition between gene-corrected cells and *TP53* mutants

We investigated how gene correction might decrease the prevalence of *TP53* mutant cells in the oral epithelium of people with FA. To run these analyses, we expanded our model to permit cells to acquire *TP53* inactivating mutations at cell division in both *FANC*^−^ (FA) and *FANC*^*+*^ (corrected) cells. Each cell division, each *FANC*^−^ and *FANC*^*+*^ cell acquires an expected number of *TP53* inactivating mutations, 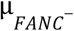 and 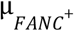, respectively. We determined the probability of *TP53* inactivation to be 7. 48 * 10^−7^ *division*^−1^ as the product of the normalized genewide *TP53* mutation rate 2. 99 * 10^−6^ *division*^−1^ (40,87) and the probability of gene inactivation conditional on mutation as determined from deep mutational scanning data (∼0.25, (88)). More information on this calculation is available in ‘*Parametrizing the proliferative advantage and mutation rate of TP53*^−^ *mutations*’. A Poisson number of inactivating mutations with 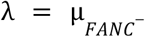 or 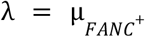 is drawn for each cell division and cells gain a fitness advantage if an inactivating mutation occurs. Acquiring multiple mutations in *TP53* did not alter clonal fitness beyond a single hit (see Discussion). *FANC*^−^ and *FANC*^*+*^ cells containing a *TP53* inactivating mutation had an additional increase in their persistence coefficient of 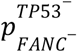 and 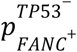, respectively. Initially, we assumed that 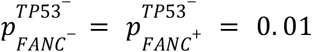, based on model fitting to the prevalence of *TP53* mutations in noncancerous tissue (see section ‘*Parametrizing the proliferative advantage and mutation rate of TP53*^−^ *mutations’* for full details).

We considered three experimental conditions in which gene correction could slow the rate at which

*TP53* mutations accumulated in tissue:

1. If *FANC*^*+*^ cells were more difficult to displace than *FANC*^−^ cells, but were otherwise identical in terms of the *TP53* mutation rate and clonal expansion speed. In this case, we examined simulation runs in which 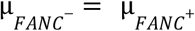 and 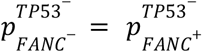
2. If *FANC*^*+*^ cells were more genetically stable than *FANC*^−^ cells and thus acquired fewer *TP53* mutations. In this case, we examined simulations in which 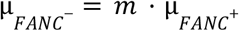 and 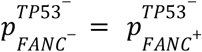, where *m* = (1. 5, 2, 4, 8).
3. If *FANC*^−^ cells permitted a faster rate of clonal expansion of *TP53* inactivating mutations than *FANC*^*+*^ cells. In this case, we examined simulations in which 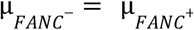 and 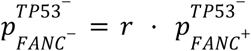, where *r* = (1. 5, 2, 4).

We conducted 300 replicates of experimental condition 1 and 100 replicates of experimental conditions 2 and 3, both with and without gene correction. Among simulations with gene correction, we examined small and large persistence coefficients for corrected cells (*p*_*corr*_ = 0. 01 and *p*_*corr*_ = 0. 1, respectively). We ran simulations on 0.33 mm^2^ (70 × 70 cells) tissue sections for up to 50 years with *k* = 30 initially corrected cells distributed with *D* = 2. Simulations without correction ran uninterrupted for the full 50 years. Simulations with correction were run for up to 50 years, but were stopped if all corrected cells were lost from the basal layer.

This stoppage criterion reflects our assumption that once all corrected cells are lost from the basal layer, the tissue behaves similarly to one never having undergone correction. To address variation in replicate numbers caused by early stoppage, mutational characteristics for samples in which gene correction was lost were sampled from a random replicate that did not undergo correction at each timepoint. We then quantified the proportion of the tissue with at least one *TP53* mutation at six month intervals for up to 46 years.

### Parametrizing the proliferative advantage and mutation rate of *TP53*^−^ mutations

To quantify how gene correction could disrupt *TP53* mutation clonal expansions in FA oral epithelium, we first parameterized *TP53*^−^ mutation rates 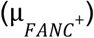 and proliferative advantages 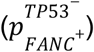 in the context of our model.

We estimated these parameters via comparison to a published study of *TP53* mutational expansion in healthy (i.e., non-FA) esophagus (48), another non-keratinized stratified squamous epithelium that is spatially-contiguous with the oral mucosa.

Martincorena et al. (48) collected 2 mm^2^ sections of esophageal tissue from study participants of different ages and deep sequenced each section to measure the number of unique *TP53* mutations and their corresponding variant allele frequencies (VAFs). Because 2 mm^2^ tissue sections are computationally expensive to simulate under the HE model, we simulated smaller tissue sections (70 × 70 cells or ∼0.33 mm^2^) and compared them to a spatially downsampled version of (48), intended to mimic results from Martincorena et al. if smaller sections were sequenced.

To approximate downsampling of the clinically-derived 2 mm^2^ samples, we first spatially reconstructed possible tissue sections that reflect the mutation data observed in each full 2 mm^2^ sample (i.e., each tissue section at a given age has *i* distinct *TP53* mutations indexed 1, …, *i*, at frequencies *f*_1_, …, *f* _*i*_). With our expected cellular density, each 2 mm^2^ tissue section is represented by a 173 × 173 cellular grid. We then place *i* mutations in cells on the grid such that each mutation is spatially contiguous and at its observed frequency, *f* _*i*_. Specifically, starting with the most frequent mutation, we choose a random starting point on the grid to assign the mutation and then mutate a random neighboring cell of the currently mutated cells to extend the spatially contiguous mutational patch until the mutation reaches its observed frequency *f* _*i*_. This process is repeated serially for each additional mutation observed in a tissue section, with the additional constraint that if the frequency of total mutations reaches 0.5 or more, the mutation that would exceed this threshold is instead assigned to an existing mutant clone background of higher frequency. After reconstructing potential tissue sections that could produce the correct number of *TP53* mutations and their VAFs in the 2 mm^2^ sections, we randomly sampled a 0.33 mm^2^ (70 × 70 cell) square subsection from each larger reconstructed sample and counted the number of mutations and their VAFs in this subsection. To align with Martincorena et al. (48), who excluded mutations sampled at frequencies below 0.0018 (108 genomic copies/(29929 cells * 2 genomic copies/cell)) in 2 mm^2^ tissue sections, we applied the equivalent threshold in our 0.33 mm^2^ sections, corresponding to a frequency 0.011 (i.e., 108 genomic copies/(4900 cells * 2 genomic copies/cell)).

To identify the *TP53* persistence coefficient value that most accurately reflects the mutation patterns observed in empirical data, we simulated 70×70 tissue sections across a range of 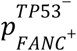 values and compared the simulated data with the downsampled empirical observations. Martincorena et al. (48) collected 844 samples from nine donors of varying ages. We examined the 560 samples from the six donors aged 20 to 55 to match the correction timescales profiled in this study. 177 samples contained detectable *TP53* mutations, which were spatially reconstructed and then downsampled as described above. Only *TP53* nonsense, missense and synonymous SNV mutations were included, with non-SNV and splice mutations excluded. Total empirical mutational prevalences (i.e., the number of unique *TP53* mutations per tissue) were computed by combining the downsampled tissue sections and the tissue sections containing no *TP53* mutations.

We conducted 100 simulation replicates for each 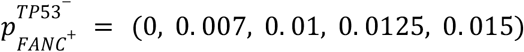 and matched two metrics between the simulated and observed data: the number of unique *TP53* mutations in the tissue and the average mutational VAF of an observed *TP53* mutation. Using the default mutation rate of the HE model resulted in an excess number of *TP53* mutations compared to empirical data, even when these mutations conferred no proliferative advantage (**Supplemental Figure 4A**). To better align the number of mutations in a tissue sample with observed data, we reasoned that the mutation rate producing mutations capable of expanding was likely lower than expected from the gene length and average mutation rate alone. To compute an adjusted mutation rate, we referenced deep mutational scanning data of *TP53* and found that approximately 25% of *TP53* mutations were expected to confer a selective benefit (88). We therefore reduced the model’s default expected number of *TP53* mutations by a factor of four, and found that this reduced mutation rate (7. 48 * 10^−7^ *division*^−1^) was capable of producing mutation counts and sizes that roughly matched Martincorena et al. (48), as shown in **Supplemental Figure 4B**.

To compare the number of distinct mutations arising, we calculated the squared differences between the mean number of mutations in the downsampled empirical data and the mean number of mutations in the simulated data for each timepoint and for each 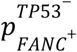 value. We then averaged these squared differences across all timepoints and identified which 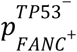 value minimized the mean squared error (**Supplemental Figure 4D**).

To find the 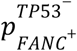 value best able to produce empirical *TP53* mutational size distributions (i.e. VAFs), we took a two step approach. First, we fit exponential distributions that described the simulated VAFs for each 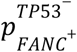 value separately at each timepoint. We fit these distributions using the R fitdist function within the fitdistrplus package (89). Before fitting, we recentered the simulated VAFs by subtracting off the lower limit of detection frequency (0.011) to better conform to the shape of the exponential distribution (i.e., bounded below at 0). Second, we then used the best fit λ at each time point and for each simulated 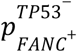 to compute the probability that an interval encompassing each empirical downsampled VAF measurement was drawn from each of the best fit distributions. Specifically, for an observed VAF measurement *f* and best fit parameter λ for a given age group and 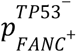, we computed the probability of the observed data as:

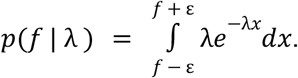

We used ε = 0. 0002 although we found that results were robust to our choice of small ε. As when fitting the distributions initially to find the best fit λs, we recentered the observed VAFs by subtracting off the lower limit of detection before evaluating the probability expressions above. We then summed the log probabilities for a given 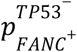 across all age groups to find the 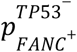 that generates simulated data that best conforms to the empirical data through time (**Supplemental Table 2, Supplemental Figure 4CE**). We did not perform exponential best fit calculations to 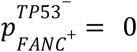 as there was only one observed mutation for the age ranges 20-23 and 24-27. Two outlier measurements diverging substantially from the overall temporal signal were excluded when determining the best fit parameters: the number of mutations in a heavy smoker aged 44-47 and the variant allele frequencies among ages 20-23 (**Supplemental Figure 4BC**).

We found that 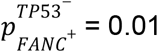 both minimized the squared error for the number of mutations and minimized the negative log-likelihood fit to the mutational VAFs, so we proceeded with 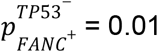 as our proliferative advantage of *TP53* mutations in healthy tissue.

Due to interactions between the *FA* and *TP53* pathways, loss of *TP53* function in cells already carrying *FANC* mutations is likely to confer a more extreme growth phenotype (i.e., 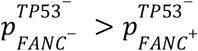, see Discussion).

To parameterize 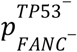, we drew on both *in vitro* and *in vivo* data from human lymphoblastoid and mouse HSC experiments (52,53) which provide consistent, quantitative evidence that *TP53* inactivation enhances proliferation to a greater degree in FA cells than non-FA cells:

1. FA and non-FA lymphoblastoid cell lines, both with and without *TP53* knockdown, were exposed to a pulse of interstrand crosslinking agents then grown in culture for 72 hours (52). In non-FA cells, *TP53* knockdown increased growth by 17%, but in FA cells, *TP53* knockdown increased growth by 40%. These values were extracted from time point H72 in Figure 3C of the original publication (52) using WebPlotDigitizer (90). This suggests 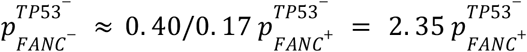.
2. HSCs from mice constructed with all combinations of *Fanca*^*+/+*^ or *Fanca*^−/–^ and *Tp53*^−/–^ or *Tp53*^+/+^ were tracked for 20 weeks (53). The *depletion* of HSCs was measured to track proliferative exhaustion due to more rapid cycling. Addition of *Tp53*^−/–^ to a *Fanca*^*+/+*^ background resulted in a decrease in abundance of 35% between weeks 8 and 20, while addition of *Tp53*^−/–^ to a *Fanca*^*-/-*^ background resulted in a decrease in abundance of 86% between weeks 8 and 20. These values were extracted from Figure 1D of the original publication (53) using WebPlotDigitizer (90). This suggests 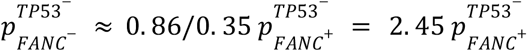, although provides less direct evidence than the growth experiments described above.

The consistency between these two studies, in different cell types from different organisms, prompted our use of an increased proliferative advantage (i.e., *r*) in the range of ∼2.4. Specifically, we tested *r* = (1. 5, 2, 4) to explore potential increases or decreases in proliferative advantage that may accompany an epithelial context.

## Code availability

The code to reproduce all analyses can be found at github.com/Colegrove/HomeostaticEpithelium and will be archived upon paper acceptance.

## References

1. Niraj J, Färkkilä A, D’Andrea AD. The Fanconi Anemia Pathway in Cancer. Annu Rev Cancer Biol. 2019 Mar 4;3(Volume 3, 2019):457–78.

2. Deans AJ, West SC. DNA interstrand crosslink repair and cancer. Nat Rev Cancer. 2011 Jul;11(7):467–80.

3. Nalepa G, Clapp DW. Fanconi anaemia and cancer: an intricate relationship. Nat Rev Cancer. 2018 Mar;18(3):168–85.

4. Johnson DE, Burtness B, Leemans CR, Lui VWY, Bauman JE, Grandis JR. Head and neck squamous cell carcinoma. Nat Rev Dis Primer. 2020 Nov 26;6(1):1–22.

5. Alter BP, Giri N, Savage SA, Rosenberg PS. Cancer in the National Cancer Institute inherited bone marrow failure syndrome cohort after fifteen years of follow-up. Haematologica. 2018 Jan 1;103(1):30–9.

6. Velleuer E, Domínguez-Hüttinger E, Rodríguez A, Harris LA, Carlberg C. Concepts of multi-level dynamical modelling: understanding mechanisms of squamous cell carcinoma development in Fanconi anemia. Front Genet [Internet]. 2023 Nov 2 [cited 2024 Sep 17];14. Available from: https://www.frontiersin.org/journals/genetics/articles/10.3389/fgene.2023.1254966/full

7. Rosenberg PS, Greene MH, Alter BP. Cancer incidence in persons with Fanconi anemia. Blood. 2003 Feb 1;101(3):822–6.

8. Carbone M, Arron ST, Beutler B, Bononi A, Cavenee W, Cleaver JE, et al. Tumour predisposition and cancer syndromes as models to study gene–environment interactions. Nat Rev Cancer. 2020 Sep;20(9):533–49.

9. Webster ALH, Sanders MA, Patel K, Dietrich R, Noonan RJ, Lach FP, et al. Genomic signature of Fanconi anaemia DNA repair pathway deficiency in cancer. Nature. 2022 Dec;612(7940):495–502.

10. van Zeeburg HJT, Snijders PJF, Wu T, Gluckman E, Soulier J, Surralles J, et al. Clinical and Molecular Characteristics of Squamous Cell Carcinomas From Fanconi Anemia Patients. JNCI J Natl Cancer Inst. 2008 Nov 19;100(22):1649–53.

11. Nguyen HT, Tang W, Webster ALH, Whiteaker JR, Chandler CM, Errazquin R, et al. Fanconi anemia-isogenic head and neck cancer cell line pairs: A basic and translational science resource. Int J Cancer. 2023;153(1):183–96.

12. Lin J, Kutler DI. Why Otolaryngologists Need to be Aware of Fanconi Anemia. Otolaryngol Clin North Am. 2013 Aug 1;46(4):567–77.

13. Kutler DI, Patel KR, Auerbach AD, Kennedy J, Lach FP, Sanborn E, et al. Natural history and management of Fanconi anemia patients with head and neck cancer: A 10-year follow-up. The Laryngoscope. 2016;126(4):870–9.

14. Dufour C, Pierri F. Modern management of Fanconi anemia. Hematology. 2022 Dec 9;2022(1):649–57.

15. Río P, Navarro S, Wang W, Sánchez-Domínguez R, Pujol RM, Segovia JC, et al. Successful engraftment of gene-corrected hematopoietic stem cells in non-conditioned patients with Fanconi anemia. Nat Med. 2019 Sep;25(9):1396–401.

16. Río P, Zubicaray J, Navarro S, Gálvez E, Sánchez-Domínguez R, Nicoletti E, et al. Haematopoietic gene therapy of non-conditioned patients with Fanconi anaemia-A: results from open-label phase 1/2 (FANCOLEN-1) and long-term clinical trials. The Lancet. 2024 Dec 21;404(10471):2584–92.

17. Siegner SM, Ugalde L, Clemens A, Garcia-Garcia L, Bueren JA, Rio P, et al. Adenine base editing efficiently restores the function of Fanconi anemia hematopoietic stem and progenitor cells. Nat Commun. 2022 Nov 12;13(1):6900.

18. Lasaga M, Río P, Vilas-Zornoza A, Planell N, Navarro S, Alignani D, et al. Gene therapy restores the transcriptional program of hematopoietic stem cells in Fanconi anemia. Haematologica. 2023 Apr 6;108(10):2652–63.

19. Río P, Navarro S, Guenechea G, Sánchez-Domínguez R, Lamana ML, Yañez R, et al. Engraftment and in vivo proliferation advantage of gene-corrected mobilized CD34+ cells from Fanconi anemia patients. Blood. 2017 Sep 28;130(13):1535–42.

20. Streichan SJ, Hoerner CR, Schneidt T, Holzer D, Hufnagel L. Spatial constraints control cell proliferation in tissues. Proc Natl Acad Sci. 2014 Apr 15;111(15):5586–91.

21. Eagle H, Levine EM. Growth Regulatory Effects of Cellular Interaction. Nature. 1967 Mar;213(5081):1102–6.

22. Lewinsohn MA, Bedford T, Müller NF, Feder AF. State-dependent evolutionary models reveal modes of solid tumour growth. Nat Ecol Evol. 2023 Apr;7(4):581–96.

23. Colom B, Alcolea MP, Piedrafita G, Hall MWJ, Wabik A, Dentro SC, et al. Spatial competition shapes the dynamic mutational landscape of normal esophageal epithelium. Nat Genet. 2020 Jun;52(6):604–14.

24. Lynch MD, Lynch CNS, Craythorne E, Liakath-Ali K, Mallipeddi R, Barker JN, et al. Spatial constraints govern competition of mutant clones in human epidermis. Nat Commun. 2017 Oct 24;8(1):1119.

25. Noble R, Burri D, Le Sueur C, Lemant J, Viossat Y, Kather JN, et al. Spatial structure governs the mode of tumour evolution. Nat Ecol Evol. 2022 Feb;6(2):207–17.

26. Abby E, Dentro SC, Hall MWJ, Fowler JC, Ong SH, Sood R, et al. Notch1 mutations drive clonal expansion in normal esophageal epithelium but impair tumor growth. Nat Genet. 2023 Feb;55(2):232–45.

27. Murai K, Dentro S, Ong SH, Sood R, Fernandez-Antoran D, Herms A, et al. p53 mutation in normal esophagus promotes multiple stages of carcinogenesis but is constrained by clonal competition. Nat Commun. 2022 Oct 20;13(1):6206.

28. Akker PC van den, Pasmooij AMG, Joenje H, Hofstra RMW, Meerman GJ te, Jonkman MF. A “late-but-fitter revertant cell” explains the high frequency of revertant mosaicism in epidermolysis bullosa. PLOS ONE. 2018 Feb 22;13(2):e0192994.

29. Jonkman MF, Scheffer H, Stulp R, Pas HH, Nijenhuis M, Heeres K, et al. Revertant Mosaicism in Epidermolysis Bullosa Caused by Mitotic Gene Conversion. Cell. 1997 Feb 21;88(4):543–51.

30. van den Akker PC, Bolling MC, Pasmooij AMG. Revertant Mosaicism in Genodermatoses: Natural Gene Therapy Right before Your Eyes. Biomedicines. 2022 Sep;10(9):2118.

31. Creighton RL, Woodrow KA. Microneedle-Mediated Vaccine Delivery to the Oral Mucosa. Adv Healthc Mater. 2019;8(4):1801180.

32. Creighton RL, Faber KA, Tobos CI, Doan MA, Guo T, Woodrow KA. Oral mucosal vaccination using integrated fiber microneedles. J Controlled Release. 2024 Mar 1;367:649–60.

33. van der Maaden K, Sekerdag E, Jiskoot W, Bouwstra J. Impact-Insertion Applicator Improves Reliability of Skin Penetration by Solid Microneedle Arrays. AAPS J. 2014 Jul 1;16(4):681–4.

34. Norman JJ, Arya JM, McClain MA, Frew PM, Meltzer MI, Prausnitz MR. Microneedle patches: Usability and acceptability for self-vaccination against influenza. Vaccine. 2014 Apr 1;32(16):1856–62.

35. Ma Y, Tao W, Krebs SJ, Sutton WF, Haigwood NL, Gill HS. Vaccine Delivery to the Oral Cavity Using Coated Microneedles Induces Systemic and Mucosal Immunity. Pharm Res. 2014 Sep 1;31(9):2393–403.

36. McNeilly CL, Crichton ML, Primiero CA, Frazer IH, Roberts MS, Kendall MAF. Microprojection arrays to immunise at mucosal surfaces. J Controlled Release. 2014 Dec 28;196:252–60.

37. Chkhaidze K, Heide T, Werner B, Williams MJ, Huang W, Caravagna G, et al. Spatially constrained tumour growth affects the patterns of clonal selection and neutral drift in cancer genomic data. PLOS Comput Biol. 2019 Jul 29;15(7):e1007243.

38. Gallaher JA, Massey SC, Hawkins-Daarud A, Noticewala SS, Rockne RC, Johnston SK, et al. From cells to tissue: How cell scale heterogeneity impacts glioblastoma growth and treatment response. PLOS Comput Biol. 2020 Feb 26;16(2):e1007672.

39. West J, Robertson-Tessi M, Anderson ARA. Agent-based methods facilitate integrative science in cancer. Trends Cell Biol. 2023 Apr 1;33(4):300–11.

40. Schenck RO, Kim E, Bravo RR, West J, Leedham S, Shibata D, et al. Homeostasis limits keratinocyte evolution. Proc Natl Acad Sci. 2022 Aug 30;119(35):e2006487119.

41. Doupé DP, Alcolea MP, Roshan A, Zhang G, Klein AM, Simons BD, et al. A Single Progenitor Population Switches Behavior to Maintain and Repair Esophageal Epithelium. Science. 2012 Aug 31;337(6098):1091–3.

42. Clayton E, Doupé DP, Klein AM, Winton DJ, Simons BD, Jones PH. A single type of progenitor cell maintains normal epidermis. Nature. 2007 Mar;446(7132):185–9.

43. Piedrafita G, Kostiou V, Wabik A, Colom B, Fernandez-Antoran D, Herms A, et al. A single-progenitor model as the unifying paradigm of epidermal and esophageal epithelial maintenance in mice. Nat Commun. 2020 Mar 18;11(1):1429.

44. Battaile KP, Bateman RL, Mortimer D, Mulcahy J, Rathbun RK, Bagby G, et al. In Vivo Selection of Wild-Type Hematopoietic Stem Cells in a Murine Model of Fanconi Anemia. Blood. 1999 Sep 15;94(6):2151–8.

45. Raya Á, Rodríguez-Pizà I, Guenechea G, Vassena R, Navarro S, Barrero MJ, et al. Disease-corrected haematopoietic progenitors from Fanconi anaemia induced pluripotent stem cells. Nature. 2009 Jul;460(7251):53–9.

46. Suzuki S, Racine RR, Manalo NA, Cantor SB, Raffel GD. Impairment of fetal hematopoietic stem cell function in the absence of Fancd2. Exp Hematol. 2017 Apr 1;48:79–86.

47. Liu S, Vivona E, Kurre P. Why hematopoietic stem cells fail in Fanconi anemia: Mechanisms and models. BioEssays. 2025;47(1):2400191.

48. Martincorena I, Fowler JC, Wabik A, Lawson ARJ, Abascal F, Hall MWJ, et al. Somatic mutant clones colonize the human esophagus with age. Science. 2018 Nov 23;362(6417):911–7.

49. Araten DJ, Golde DW, Zhang RH, Thaler HT, Gargiulo L, Notaro R, et al. A Quantitative Measurement of the Human Somatic Mutation Rate. Cancer Res. 2005 Sep 15;65(18):8111–7.

50. Sala-Trepat M, Boyse J, Richard P, Papadopoulo D, Moustacchi E. Frequencies of HPRT− lymphocytes and glycophorin A variants erythrocytes in Fanconi anemia patients, their parents and control donors. Mutat Res Mol Mech Mutagen. 1993 Sep 1;289(1):115–26.

51. Evdokimova VN, McLoughlin RK, Wenger SL, Grant SG. Use of the glycophorin A somatic mutation assay for rapid, unambiguous identification of Fanconi anemia homozygotes regardless of GPA genotype. Am J Med Genet A. 2005;135A(1):59–65.

52. Ceccaldi R, Parmar K, Mouly E, Delord M, Kim JM, Regairaz M, et al. Bone Marrow Failure in Fanconi Anemia Is Triggered by an Exacerbated p53/p21 DNA Damage Response that Impairs Hematopoietic Stem and Progenitor Cells. Cell Stem Cell. 2012 Jul 6;11(1):36–49.

53. Li X, Wilson AF, D. W, Pang Q. Cell-Cycle-Specific Function of p53 in Fanconi Anemia Hematopoietic Stem and Progenitor Cell Proliferation. Stem Cell Rep. 2018 Feb 13;10(2):339–46.

54. Freie B, Li X, Ciccone SLM, Nawa K, Cooper S, Vogelweid C, et al. Fanconi anemia type C and p53 cooperate in apoptosis and tumorigenesis. Blood. 2003 Dec 1;102(12):4146–52.

55. Houghtaling S, Granville L, Akkari Y, Torimaru Y, Olson S, Finegold M, et al. Heterozygosity for p53 (Trp53+/−) Accelerates Epithelial Tumor Formation in Fanconi Anemia Complementation Group D2 (Fancd2) Knockout Mice. Cancer Res. 2005 Jan 21;65(1):85–91.

56. Armitage P, Doll R. The Age Distribution of Cancer and a Multi-stage Theory of Carcinogenesis. Br J Cancer. 1954 Mar;8(1):1–12.

57. Durrett R. Branching Process Models of Cancer. In: Durrett R, editor. Branching Process Models of Cancer [Internet]. Cham: Springer International Publishing; 2015 [cited 2025 Jun 17]. p. 1–63. Available from: 10.1007/978-3-319-16065-8_1

58. Colom B, Herms A, Hall MWJ, Dentro SC, King C, Sood RK, et al. Mutant clones in normal epithelium outcompete and eliminate emerging tumours. Nature. 2021 Oct;598(7881):510–4.

59. Murai K, Skrupskelyte G, Piedrafita G, Hall M, Kostiou V, Ong SH, et al. Epidermal Tissue Adapts to Restrain Progenitors Carrying Clonal p53 Mutations. Cell Stem Cell. 2018 Nov 1;23(5):687-699.e8.

60. Alcolea MP, Greulich P, Wabik A, Frede J, Simons BD, Jones PH. Differentiation imbalance in single oesophageal progenitor cells causes clonal immortalization and field change. Nat Cell Biol. 2014 Jun;16(6):612–9.

61. Choate KA, Lu Y, Zhou J, Choi M, Elias PM, Farhi A, et al. Mitotic Recombination in Patients with Ichthyosis Causes Reversion of Dominant Mutations in KRT10. Science. 2010 Oct;330(6000):94–7.

62. Jones KB, Furukawa S, Marangoni P, Ma H, Pinkard H, D’Urso R, et al. Quantitative Clonal Analysis and Single-Cell Transcriptomics Reveal Division Kinetics, Hierarchy, and Fate of Oral Epithelial Progenitor Cells. Cell Stem Cell. 2019 Jan 3;24(1):183-192.e8.

63. Sender R, Milo R. The distribution of cellular turnover in the human body. Nat Med. 2021 Jan;27(1):45–8.

64. Pan Q, Nicholson AM, Barr H, Harrison L, Wilson GD, Burkert J, et al. Identification of Lineage-Uncommitted, Long-Lived, Label-Retaining Cells in Healthy Human Esophagus and Stomach, and in Metaplastic Esophagus. Gastroenterology. 2013 Apr 1;144(4):761–70.

65. Renaudin X, Rosselli F. The FANC/BRCA Pathway Releases Replication Blockades by Eliminating DNA Interstrand Cross-Links. Genes. 2020 May;11(5):585.

66. Rageul J, Kim H. Fanconi anemia and the underlying causes of genomic instability. Environ Mol Mutagen. 2020;61(7):693–708.

67. Taylor AMR, Rothblum-Oviatt C, Ellis NA, Hickson ID, Meyer S, Crawford TO, et al. Chromosome instability syndromes. Nat Rev Dis Primer. 2019 Sep 19;5(1):1–20.

68. Hughes AD, Kurre P. The impact of clonal diversity and mosaicism on haematopoietic function in Fanconi anaemia. Br J Haematol. 2022;196(2):274–87.

69. Waisfisz Q, Morgan NV, Savino M, de Winter JP, van Berkel CGM, Hoatlin ME, et al. Spontaneous functional correction of homozygous Fanconi anaemia alleles reveals novel mechanistic basis for reverse mosaicism. Nat Genet. 1999 Aug;22(4):379–83.

70. Persico I, Fiscarelli I, Pelle A, Faleschini M, Pasini B, Savoia A, et al. Phenotype reversion as “natural gene therapy” in Fanconi anemia by a gene conversion event. Front Genet [Internet]. 2023 Sep 18 [cited 2025 Feb 14];14. Available from: https://www.frontiersin.org/journals/genetics/articles/10.3389/fgene.2023.1240758/full

71. Soulier J, Leblanc T, Larghero J, Dastot H, Shimamura A, Guardiola P, et al. Detection of somatic mosaicism and classification of Fanconi anemia patients by analysis of the FA/BRCA pathway. Blood. 2005 Feb 1;105(3):1329–36.

72. Ten Foe JRL, Kwee ML, Rooimans MA, Oostra AB, Veerman AJP, van Weel M, et al. Somatic Mosaicism in Fanconi Anemia: Molecular Basis and Clinical Significance. Eur J Hum Genet. 1997 May;5(3):137–48.

73. Gross M, Hanenberg H, Lobitz S, Friedl R, Herterich S, Dietrich R, et al. Reverse mosaicism in Fanconi anemia: natural gene therapy via molecular self-correction. Cytogenet Cell Genet. 2003 Apr 14;98(2–3):126–35.

74. Asur RS, Kimble DC, Lach FP, Jung M, Donovan FX, Kamat A, et al. Somatic mosaicism of an intragenic duplication in both fibroblast and peripheral blood cells observed in a Fanconi anemia patient leads to milder phenotype. Mol Genet Genomic Med. 2018;6(1):77–91.

75. Errazquin R, Page A, Suñol A, Segrelles C, Carrasco E, Peral J, et al. Development of a mouse model for spontaneous oral squamous cell carcinoma in Fanconi anemia. Oral Oncol. 2022 Nov 1;134:106184.

76. Jaber S, Toufektchan E, Lejour V, Bardot B, Toledo F. p53 downregulates the Fanconi anaemia DNA repair pathway. Nat Commun. 2016 Apr 1;7(1):11091.

77. Peake JD, Horne KI, Noguchi C, Gilligan JP, Noguchi E. The p53 DNA damage response and Fanconi anemia DNA repair pathway protect against acetaldehyde-induced replication stress in esophageal keratinocytes. Cell Cycle. 2023 Sep 17;22(18):2088–96.

78. Piipponen M, Riihilä P, Nissinen L, Kähäri VM. The Role of p53 in Progression of Cutaneous Squamous Cell Carcinoma. Cancers. 2021 Jan;13(18):4507.

79. Kim MP, Lozano G. Mutant p53 partners in crime. Cell Death Differ. 2018 Jan;25(1):161–8.

80. Wang M, Brandt LTL, Wang X, Russell H, Mitchell E, Kamimae-Lanning AN, et al. Genotoxic aldehyde stress prematurely ages hematopoietic stem cells in a p53-driven manner. Mol Cell. 2023 Jul 20;83(14):2417-2433.e7.

81. Tsuchida CA, Wasko KM, Hamilton JR, Doudna JA. Targeted nonviral delivery of genome editors in vivo. Proc Natl Acad Sci. 2024 Mar 12;121(11):e2307796121.

82. Saha K, Sontheimer EJ, Brooks PJ, Dwinell MR, Gersbach CA, Liu DR, et al. The NIH Somatic Cell Genome Editing program. Nature. 2021 Apr;592(7853):195–204.

83. Subramaniam S, Akay M, Anastasio MA, Bailey V, Boas D, Bonato P, et al. Grand Challenges at the Interface of Engineering and Medicine. IEEE Open J Eng Med Biol. 2024;5:1–13.

84. Musunuru K, Grandinette SA, Wang X, Hudson TR, Briseno K, Berry AM, et al. Patient-Specific In Vivo Gene Editing to Treat a Rare Genetic Disease. N Engl J Med. 2025 Jun 11;392(22):2235–43.

85. Bravo RR, Baratchart E, West J, Schenck RO, Miller AK, Gallaher J, et al. Hybrid Automata Library: A flexible platform for hybrid modeling with real-time visualization. PLOS Comput Biol. 2020 Mar 10;16(3):e1007635.

86. Simons BD. Deep sequencing as a probe of normal stem cell fate and preneoplasia in human epidermis. Proc Natl Acad Sci. 2016 Jan 5;113(1):128–33.

87. Schenck RO, Brosula G, West J, Leedham S, Shibata D, Anderson ARA. Gattaca: Base-Pair Resolution Mutation Tracking for Somatic Evolution Studies using Agent-based Models. Mol Biol Evol. 2022 Apr 1;39(4):msac058.

88. Kotler E, Shani O, Goldfeld G, Lotan-Pompan M, Tarcic O, Gershoni A, et al. A Systematic p53 Mutation Library Links Differential Functional Impact to Cancer Mutation Pattern and Evolutionary Conservation. Mol Cell. 2018 Jul 5;71(1):178-190.e8.

89. Delignette-Muller ML, Dutang C. fitdistrplus: An R Package for Fitting Distributions. J Stat Softw. 2015 Mar 20;64:1–34.

90. Rohatgi A. WebPlotDigitizer: Version 5.2 [Internet]. 2024. Available from: https://automeris.io/

91. Guerra L, Diociaiuti A, El Hachem M, Castiglia D, Zambruno G. Ichthyosis with confetti: clinics, molecular genetics and management. Orphanet J Rare Dis. 2015 Sep 17;10(1):115.

92. Nomura T, Suzuki S, Miyauchi T, Takeda M, Shinkuma S, Fujita Y, et al. Chromosomal inversions as a hidden disease-modifying factor for somatic recombination phenotypes. JCI Insight [Internet]. 2018 Mar 22 [cited 2025 Jun 23];3(6). Available from: https://insight.jci.org/articles/view/97595

93. Lim YH, Qiu J, Saraceni C, Burrall BA, Choate KA. Genetic Reversion via Mitotic Recombination in Ichthyosis with Confetti due to a KRT10 Polyalanine Frameshift Mutation. J Invest Dermatol. 2016 Aug 1;136(8):1725–8.

94. Burger B, Spoerri I, Schubert M, Has C, Itin PH. Description of the natural course and clinical manifestations of ichthyosis with confetti caused by a novel KRT10 mutation. Br J Dermatol. 2012 Feb 1;166(2):434–9.

