## Supplemental File for "Epithelial competition determines gene therapy potential to suppress Fanconi Anemia oral cancer risk"

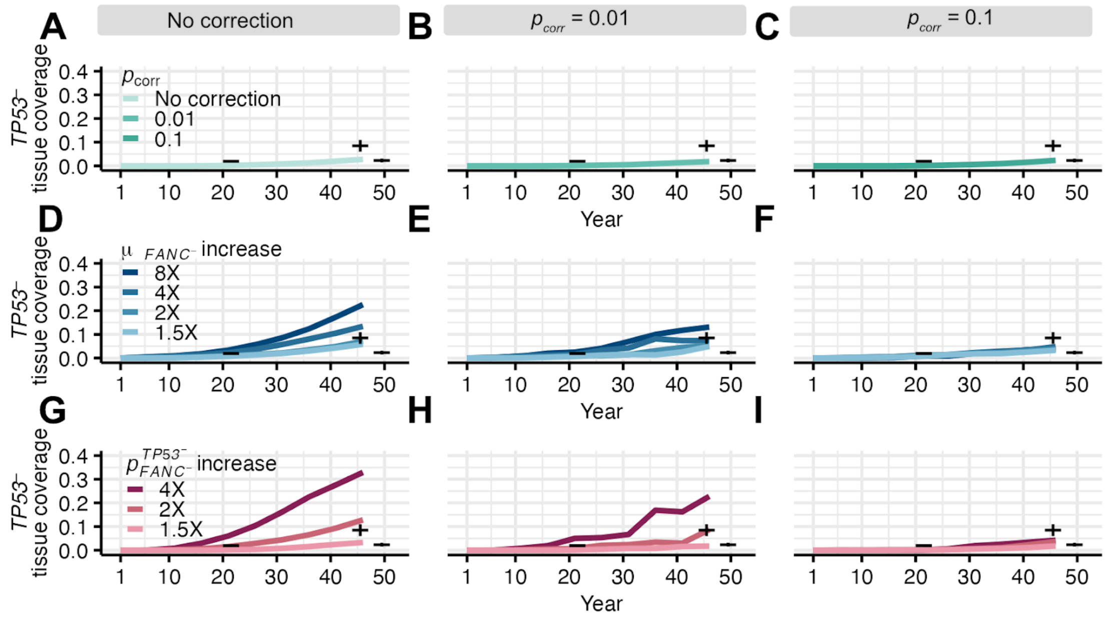


**Supplemental Figure 1: Gene correction reduces spread of *TP53* mutations through FA tissue over time.**

**(A-I)** *TP53*⁻ tissue coverage tracked over time for 46 years on 0.33 $mm^2$ simulated tissue sections with or without single gene correction ($k = 30$ cells and $D = 2$). Columns represent conditions with no correction, $p_{corr} = 0.01$ and $p_{corr} = 0.1$. Rows represent the experimental conditions as described in Figure 4. Black crosses represent *TP53* tissue coverage ranges from Martincorena et al. (48) normal esophagus, with vertical lines indicating the observed range. 300 simulations were performed for each condition within each persistence coefficient for **A-C** and 100 simulations were performed for **D-I**.

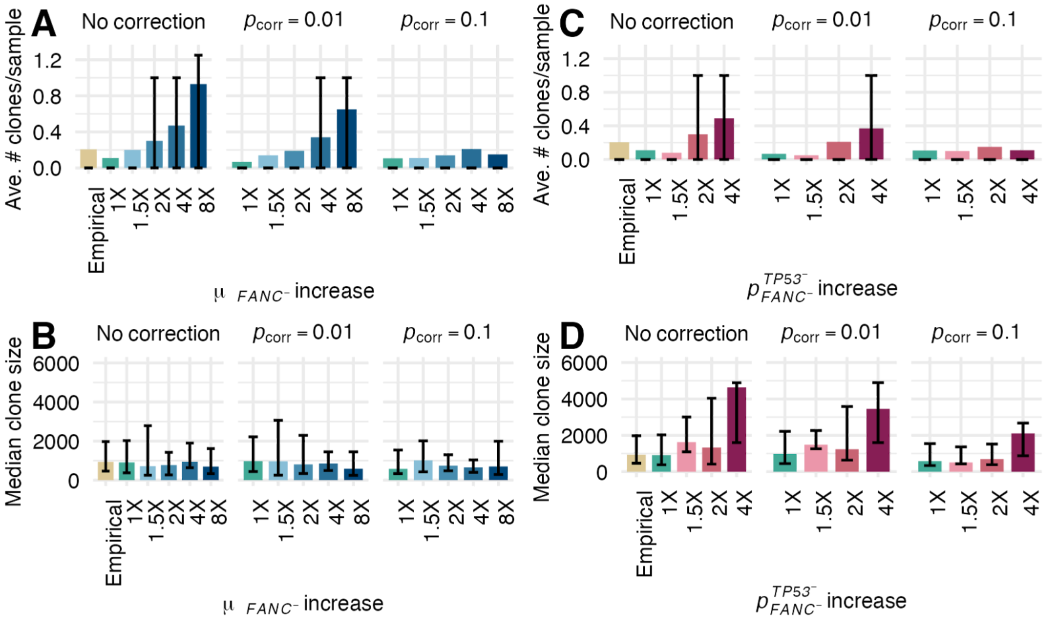


**Supplemental Figure 2: Elevated mutation rates increase the number of *TP53⁻* clones, while increased persistence coefficients increase *TP53⁻* clone sizes.**

**(A-D)** The average number and size of *TP53⁻* clones after 46 years on 0.33 $mm^2$ simulated tissue sections with ($p_{corr} = 0.01$ or $0.1$) or without gene correction. The 1X condition in each panel (green) represents simulations where *TP53* mutation rates and persistence coefficients are equal across *FANC⁻* and *FANC⁺* cells. The empirical condition in each panel (yellow) represents data from downsampled normal esophageal tissue in ages 36-55 from Martincorena et al. (48). 300 tissue simulations of the 1X condition and 100 tissue simulations of all other conditions were performed. Bars indicate interquartile ranges.

**(A)** Average number of distinct *TP53⁻* clones when the *TP53* mutation rate is elevated in *FANC⁻* cells ($\mu_{FANC^-} = m \cdot \mu_{FANC^+}$, $m = (1.5, 2, 4, 8)$). **(B)** Median *TP53⁻* clone size under the same conditions as **(A)**.

**(C)** Average number of distinct *TP53⁻* clones when *TP53* persistence coefficients are elevated in *FANC⁻* cells ($p^{TP53^-}_{FANC^-} = r \cdot p^{TP53^-}_{FANC^+}$, $r = (1.5, 2, 4)$). **(D)**. Median *TP53⁻* clone size under the same conditions as **(C)**.

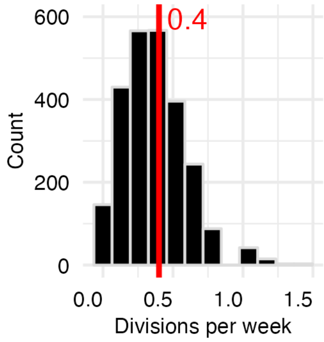


**Supplemental Figure 3: Basal cell division rates in the default HomeostaticEpidermis model.** Distribution of average cell division rates across basal layer positions in the HomeostaticEpidermis model over 100 model timesteps on a 50x50 cell grid. The red line indicates the mean division rate (0.4 divisions per week).

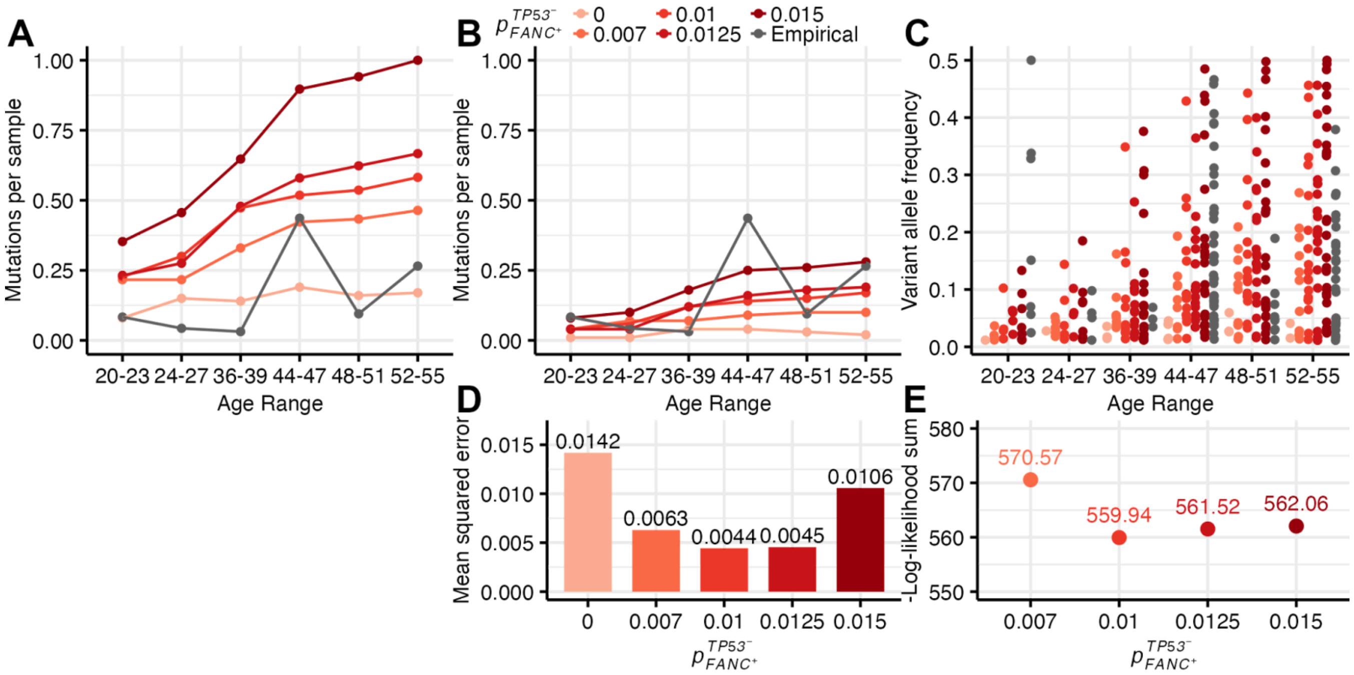


**Supplemental Figure 4: Parameterizing the proliferative advantage and mutation rate of *TP53* mutations**

**(A)** The number of *TP53* mutations per 0.33 mm$^2$ tissue sample in simulations with varying persistence coefficients ($p^{TP53^-}_{FANC^+}$ = 0, 0.007, 0.01, 0.0125, 0.015) using the default mutation rate of the HomeostaticEpidermis model (colored), compared to empirical data (grey). 100 tissue simulations were performed for each age range and persistence coefficient. **(B)** The number of *TP53* mutations per sample in simulations with varying persistence coefficients using the calibrated mutation rate ($\mu_{FANC^+}$). **(C)** Variant allele frequencies of mutation meeting the lower limit of detection in simulated (colored) and empirical (grey) tissue. **(D)** Mean squared error between the simulated and empirical data for the number of *TP53* mutations across various $p^{TP53^-}_{FANC^+}$ values. Age range 44-47 was excluded from the mean squared error computation. **(E)** Negative log-likelihood comparisons of empirical variant allele frequencies against simulated distributions across $p^{TP53^-}_{FANC^+}$ values. Age range 20-23 was excluded from negative log-likelihood computation.

**Supplemental Table 1: Biological contextualization of persistence coefficient values ($p_{corr}$)**

| $p_{corr}$ | Interpretation | Approximate clone size distribution in adulthood |
|---|---|---|
| 0 | Corrected cells lack a competitive advantage and expand by neutral drift, with clone sizes similar to synonymous mutations. | 5 clones detected across 100 0.33mm$^2$ simulated sections between ages 44-55. Clone sizes span 0.01-0.02 mm$^2$ within the interquartile range[b] |
| 0.01 | Corrected cells confer a competitive advantage to expand at rates comparable to *TP53* mutations in normal esophagus. | 76 clones were detected across 287 downsampled 0.33mm$^2$ sections between ages 44-55. Clone sizes span 0.03-0.12 mm$^2$ within the interquartile range[a] |
| 0.02 | Corrected cells confer a competitive advantage to expand at rates comparable to *NOTCH1* mutations in normal esophagus. | 307 clones were detected across 287 downsampled 0.33mm$^2$ sections between ages 44-55. Clones sizes span 0.06-0.19 mm$^2$ within the interquartile range[a] |
| 1.0 | Corrected cells confer a competitive advantage to expand at the maximum possible rate in our model. | Clones fully saturate 0.33 mm$^2$ simulated tissue sections by 3 years in 100% of cases[b] |
| - | Degree of expansion in ichthyosis with confetti revertant clones | By ages 18-42, individuals typically developed hundreds of 2-10 mm diameter revertant patches, with capability to exceed 4 cm (30,61,91–94). |

[a] Clone sizes based on downsampled tissue sections from Martincorena et al. (48)
[b] Clone sizes based on oral epithelial model simulation estimates

**Supplemental Table 2**: **Best-fit exponential parameters and corresponding negative log-likelihood values across $p_{FANC^+}^{TP53^-}$ conditions and age ranges**

| | | $p_{FANC^+}^{TP53^-} = 0.007$ | | $p_{FANC^+}^{TP53^-} = 0.01$ | | $p_{FANC^+}^{TP53^-} = 0.0125$ | | $p_{FANC^+}^{TP53^-} = 0.015$ | |
|---|---|---|---|---|---|---|---|---|---|
| Age range | # of obs | λ | $p(\bar{f}_{obs}\|\lambda)$ | λ | $p(\bar{f}_{obs}\|\lambda)$ | λ | $p(\bar{f}_{obs}\|\lambda)$ | λ | $p(\bar{f}_{obs}\|\lambda)$ |
| 24-27 | 4 | 47.87 | 24.21 | 23.78 | 22.79 | 20.96 | 22.8 | 23.43 | 22.79 |
| 36-39 | 3 | 17.77 | 16.96 | 13.59 | 17.26 | 15.55 | 17.09 | 10.29 | 17.7 |
| 44-47 | 41 | 13.59 | 300.35 | 8.34 | 286.93 | 8.67 | 287.44 | 7.31 | 285.78 |
| 48-51 | 9 | 11.56 | 53.9 | 6.31 | 56.85 | 6.4 | 56.76 | 5.68 | 57.49 |
| 52-55 | 26 | 9.21 | 175.15 | 5.93 | 176.11 | 5.07 | 177.44 | 4.67 | 178.29 |
| **Total** | | | **570.57** | | **559.94** | | **561.53** | | **562.05** |
